# *Aspergillus fumigatus* SidF is a dual substrate acyltransferase involved in biosynthesis of both fusarinine- and ferrichrome-type siderophores

**DOI:** 10.1101/2024.08.20.608788

**Authors:** Patricia Caballero, Annie Yap, Simon Oberegger, Isidor Happacher, Thanalai Poonsiri, Stefano Benini, Hubertus Haas

## Abstract

The human pathogen *Aspergillus fumigatus* produces fusarinine-type (FusTS) and ferrichrome-type siderophores (FchTS), both of which have been shown to be crucial for virulence of this mold. After the common first siderophore biosynthetic step, SidA-catalyzed hydroxylation of ornithine, the pathway splits. For FusTS biosynthesis, SidF incorporates an anhydromevalonyl group, while for FchTS biosynthesis, SidL and an as yet unknown enzyme incorporate an acetyl group. The transacylases SidF and SidL share only limited similarity in their C-terminal GNAT (Gcn5-related N-acetyltransferases) motif-containing domains. SidF is transcriptionally induced by iron limitation and localizes to peroxisomes, whereas SidL is a cytosolic enzyme with largely iron-independent expression.

Here, we discovered that simultaneous inactivation of both SidF and SidL abolished the biosynthesis of both FusTS and FchTS and caused a growth defect under iron limitation, similar to the inactivation of SidA. Biosynthesis of both FusTS and FchTS depended on both the unique N-terminal and the GNAT motif-containing C-terminal SidF domains. In conclusion, SidF is the hitherto unknown FchTS biosynthetic enzyme: in contrast to SidL, SidF is a bifunctional enzyme accepting acetyl-CoA and anhydromevalonyl-CoA as substrates for biosynthesis of both FusTS and FchTS. Furthermore, this study revealed interdependence of FusTS and FchTS production, and that the peroxisomal localization of FusTS enzymes is important for optimizing FusTS production at the expense of FchTS. Phylogenetic analyses supported the relevance of these findings to other fungal species and revealed overlapping but distinct consensus sequences for the GNAT motifs of SidL and SidF, most likely reflecting their different substrate specificities.

**IMPORTANCE:** Adaptation to the host niche is key for any pathogenic organism. *Aspergillus fumigatus* is a major fungal pathogen causing 90% of invasive aspergillosis cases, which is associated with a high mortality rate. Siderophore-mediated iron acquisition has been shown to be essential for virulence of *A. fumigatus* and other fungal pathogens. In recent years, the hyphal siderophore biosynthetic pathway has been largely elucidated with exception of a single unknown enzyme, which we identified here as SidF. In contrast to another siderophore biosynthetic acyltransferase, SidL, SidF is a bifunctional enzyme accepting different substrates. As simultaneous inactivation of SidF and SidL, which share a common protein domain and a common substrate, blocks the biosynthesis of all siderophores, simultaneous targeting of SidF and SidL may allow development of new antifungal drugs. Phylogenetic analyses supported the relevance of these findings to other fungal species Moreover, this study clarified the rational for partial peroxisomal localization of siderophore biosynthesis and their metabolic interdependence.

Here, we discovered that simultaneous inactivation of both SidF and SidL abolished the biosynthesis of both FusTS and FchTS and caused a growth defect under iron limitation, similar to the inactivation of SidA. Biosynthesis of both FusTS and FchTS depended on both the unique N-terminal and the GNAT motif-containing C-terminal SidF domains. Taken together, SidF is the hitherto unknown FchTS biosynthetic enzyme: in contrast to SidL, SidF is a bifunctional enzyme accepting acetyl-CoA and anhydromevalonyl-CoA as substrates for biosynthesis of both FusTS and FchTS. Moreover, this study revealed interdependence of FusTS and FchTS production, and that peroxisomal localization of FusTS enzymes is important for optimizing FusTS production at the expense of FchTS.

## INTRODUCTION

The saprobic fungus *Aspergillus fumigatus* is one of the most common opportunistic fungal pathogens causing life-threatening invasive infectious diseases, particularly in immunocompromised patients, known as aspergillosis (1,2). Among many factors, iron homeostasis is known to be essential for fungal survival in both the host and the environment, and is key to the virulence of *A. fumigatus* in various infection models (3). *Aspergillus fumigatus* employs two high-affinity iron uptake systems: reductive iron assimilation (RIA) and siderophore-mediated iron acquisition (SIA) (3). RIA starts with extracellular reduction of ferric iron to ferrous iron by plasma membrane-localized metalloreductases, such as FreB (4), followed by the re-oxidation to ferric iron by the iron oxidase FetC and uptake by the ferric iron permease FtrA (5). Siderophores are low-molecular-mass, ferric iron-chelating molecules. *A. fumigatus* synthesizes four hydroxamate-class siderophores: two fusarinine-type (FusTS), fusarinine C and triacetylfusarinine C (TAFC), and two ferrichrome-type siderophores (FchTS), ferricrocin (FC) and hydroxyferricrocin. Fusarinine C, TAFC, and FC are secreted by *A. fumigatus* in order to capture iron from the surrounding environment (6,7). Additionally, FC is employed for the intracellular handling of iron within hyphae, while hydroxyferricrocin is used for the storage of iron in conidia (7–9). A scheme of the siderophore biosynthetic pathway is shown in Figure 1. The initial committed step in the biosynthesis of all four siderophores is the formation of N^5^-hydroxyornithine from ornithine, which is catalyzed by the monooxygenase SidA (5). Then, the pathways for the biosynthesis of FusTS and FchTS diverge due to the incorporation of different acyl groups. For the biosynthesis of FchTS, N^5^-hydroxyornithine is acetylated to N^5^-acetyl-N^5^-hydroxyornithine by the transacetylase SidL, which is constitutively active, whereas an as yet unidentified enzyme complements this function during iron starvation (7). Subsequently, the non-ribosomal peptide synthetase (NRPS) SidC assembles FC from three N^5^-acetyl-N^5^-hydroxyornithine, two glycine, and one serine residues (7). The hydroxylation of FC by an as yet-unknown enzyme results in the formation of hydroxiferricrocin (10,11). For FusTS biosynthesis, the transacylase SidF uses anhydromevalonyl-CoA for formation of N^5^-anhydromevalonyl-N^5^-hydroxyornithine (7). Anhydromevalonyl-CoA is derived from mevalonate involving the mevalonyl-CoA ligase SidI and the mevalonyl-CoA hydratase SidH (12). The linkage of three N^5^-anhydromevalonyl-N^5^-hydroxyornithines by the NRPS SidD results in the formation of fusarinine C and its triple N^2^-acetylation, catalyzed by SidG, results in TAFC (7). Following extracellular chelation, ferric siderophores are internalized by the cell via specific siderophore iron transporters: Sit1 and Sit2 for FchTS such as FC, MirD for fusarinine C, and MirB for TAFC (13–16). For intracellular release of iron, ferric fusarinine C and TAFC are hydrolyzed by the esterases SidJ and EstB, respectively (17–19). Three enzymes involved in FusTS biosynthesis have been shown to localize to peroxisomes, namely peroxisomal targeting sequence (PTS) 1-containing SidH and SidF and PTS2-containing SidI (20). In contrast, the acyltransferase SidL has been demonstrated to localize to the cytosol (11). The mislocalization of one or two of the enzymes responsible for TAFC production to the cytosol has been shown to impair TAFC production in both *A. fumigatus* and *A. nidulans* (20). In contrast to RIA, SIA has been demonstrated to be crucial for virulence, as the lack of siderophore biosynthesis (lack of SidA) resulted in avirulence, whereas the absence of either FusTS (lack of SidI, SidH, SidF, or SidD) or FchTS (lack of SidC), or TAFC uptake (lack of MirB) attenuated virulence in several infection models (3,5,7,12,13,21,22).

**Figure 1:**
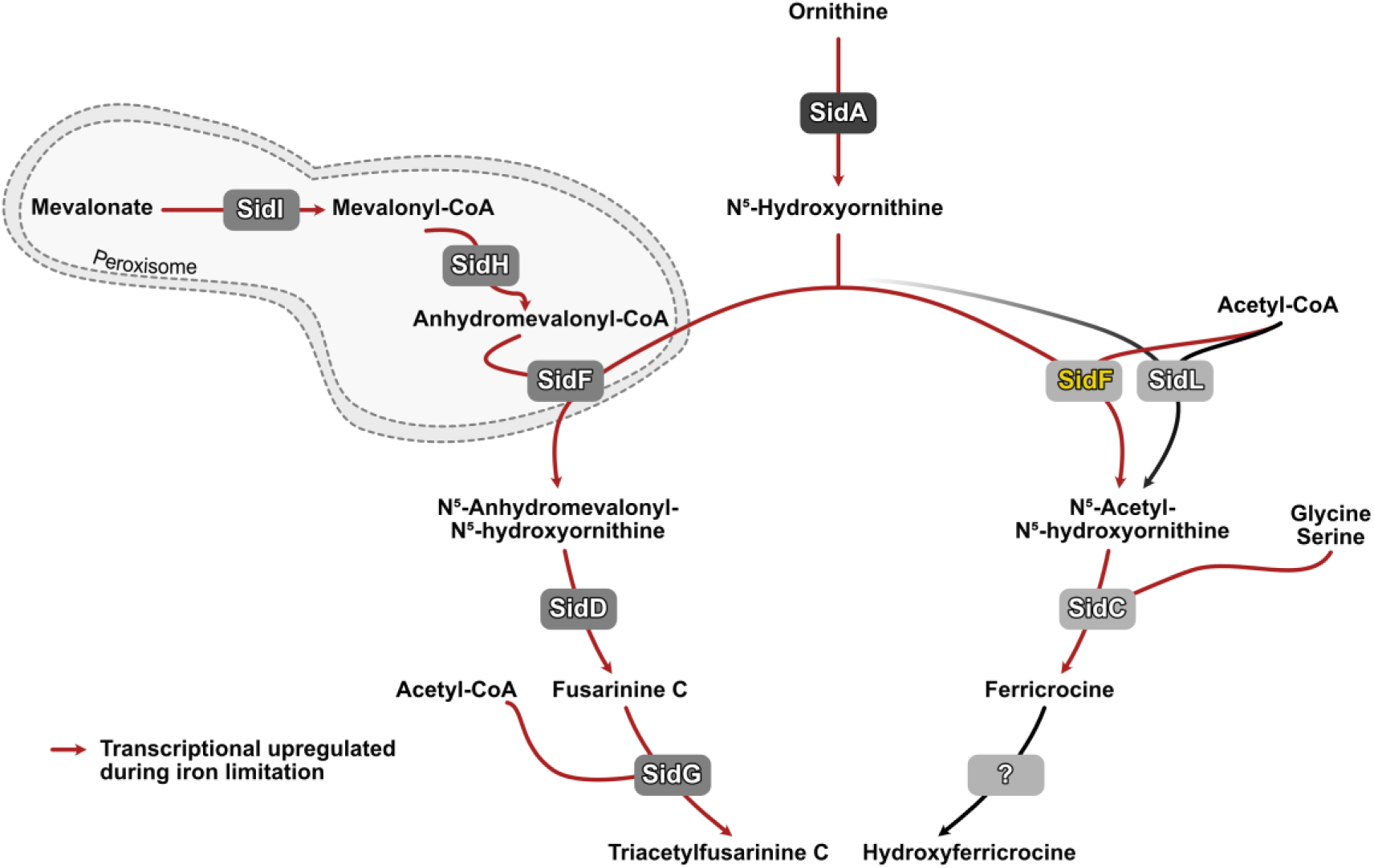
The *A. fumigatus* siderophore biosynthetic pathway. Scheme was adapted from (3,52). Red arrows indicate transcriptional upregulation during iron limitation. The SidF function identified in this study is highlighted in yellow.

Iron homeostasis in *A. fumigatus* depends on two iron-sensing transcription factors, SreA and HapX (23–25). During iron limitation, HapX represses iron-consuming pathways to spare iron and induces iron uptake via SIA and RIA (26). Iron excess converts HapX into an activator of vacuolar iron deposition mediating iron resistance (27). In contrast, SreA represses SIA and RIA during iron sufficiency to avoid toxicity caused by excessive iron uptake (28). HapX, but not SreA, was shown to be critical for virulence of *A. fumigatus*, highlighting the importance of efficient adaptation to iron limitation for fungal pathogenicity (26).

Recently, *in vitro* biochemical studies indicated that the acyltransferase SidF accepts not only anhydromevalonyl-CoA but also acetyl-CoA as a substrate, like SidL (29). Both SidF and SidL belong to the GNAT (Gcn5-related N-acetyltransferases) family of proteins, which share limited similarity only in the C-terminal GNAT motif-containing domain (11) and have significant differences: SidF is transcriptionally repressed by iron via SreA and localized to peroxisomes (11,20,28), whereas SidL is a cytosolic enzyme with largely iron-independent expression (11). Here we investigated the possible involvement of SidF in the biosynthesis of FchTS, the link between FusTS and FchTS production and the role of peroxisomes in siderophore production in *A. fumigatus*.

## RESULTS

### The acetyltransferase SidF is involved in FchTS biosynthesis

To investigate the potential involvement of SidF in the biosynthesis of FchTS, we generated mutants lacking the genes encoding SidF (*ΔsidF*), SidL (*ΔsidL*) or both acyltransferases (*ΔsidFΔsidL*) in *A. fumigatus s*train A1160+, termed wild type (wt) here, which is derived from the clinical isolate CEA10 (30). In a next step, the growth pattern of the generated mutant strains was compared to wt and to a previously generated mutant in the same genetic background lacking SidA (*ΔsidA*) and consequently biosynthesis of all siderophores (13) on solid medium reflecting different iron availability (Figure 2). Largely similar to wt, mutants lacking either SidF (*ΔsidF*) or SidL (*ΔsidL*) were able to grow under iron limitation, i.e., without iron supplementation or with 1 µM iron supplementation, even in the presence of the ferrous iron-specific chelator bathophenanthrolinedisulfonic acid (BPS), which inhibits siderophore-independent high-affinity iron acquisition mediated by RIA (5). This can be explained by the fact that even in the absence of RIA, *ΔsidF* is able to acquire iron via FC secretion and that SidL can utilize FusTS-mediated iron uptake (6). Compared to wt, lack of SidF (*ΔsidF*) slightly decreased radial growth under iron limitation and lack of SidL (*ΔsidL*) decreased conidiation during iron limitation, which is most likely explained by the role of FC in intracellular iron distribution for conidiation (8). In contrast to the single mutants, the mutant lacking both acyltransferases (*ΔsidLΔsidF*) showed a strong growth and conidiation defect, particularly during iron limitation, which was rescued by high iron supplementation (10 mM), similar to *ΔsidA.* Interestingly, the *ΔsidLΔsidF* mutant exhibited an even more pronounced growth defect at 1 µM iron compared to *ΔsidA*.

**Figure 2:**
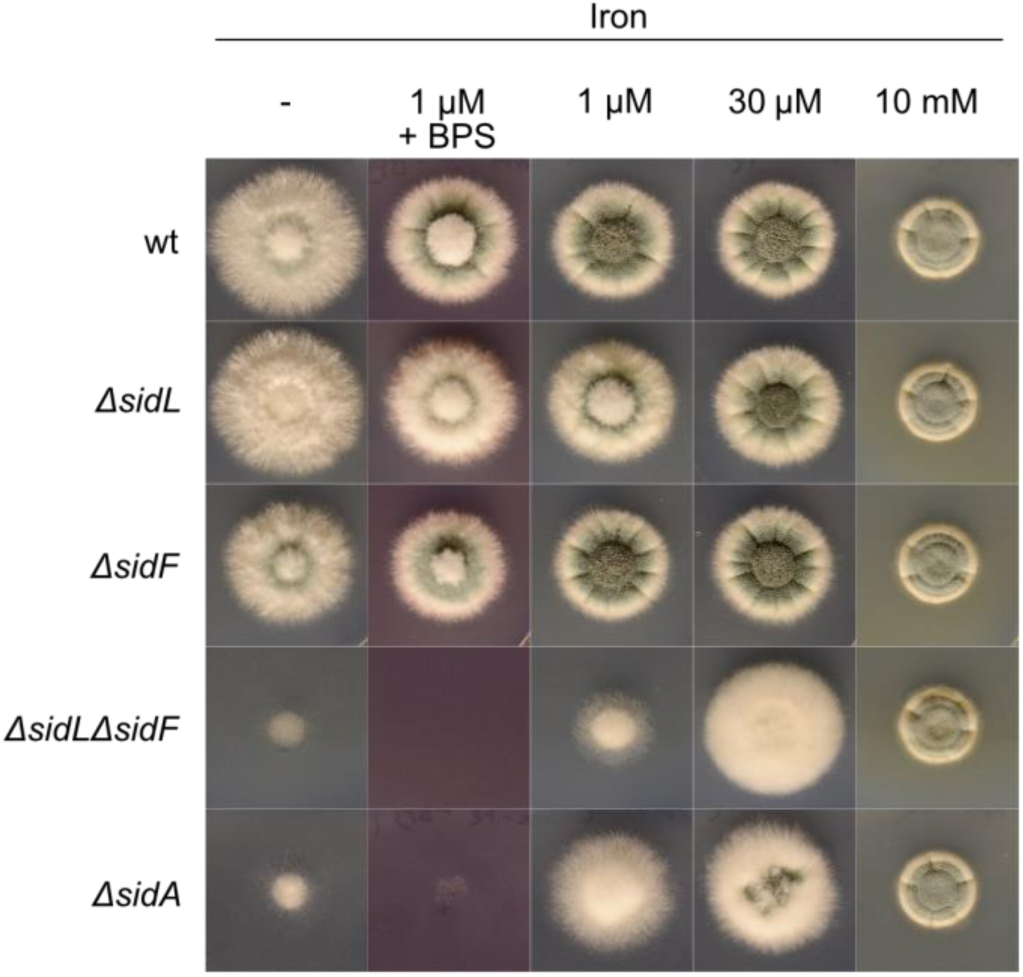
Simultaneous inactivation of SidL and SidF largely phenocopies lack of SidA, i.e. causes a growth defect under iron limitation. *Aspergillus fumigatus* wt and *ΔsidL*, *ΔsidF*, *ΔsidA* and *ΔsidLΔsidF* mutant strains were point-inoculated using 10^4^ spores on minimal medium without iron addition (-) or supplemented with FeSO_4_ to final concentrations of 1 µM, 30 µM or 10 mM. The ferrous iron-specific BPS was used in a final concentration of 200 µM. Plates were incubated at 37 °C for 48 h. Conidiation is reflected by greenish coloration of the fungal colonies due to the conidial DHN-melanin (53).

In liquid shake flask cultures, *ΔsidF* and *ΔsidL* mutant strains showed largely wt-like growth under all conditions tested, i.e., media without iron (-Fe), 1 µM iron supplementation (1 µM Fe) and 30 µM (+Fe) iron supplementation (Figure 3). In contrast, *ΔsidFΔsidL* and *ΔsidA* were not able to grow under -Fe conditions and showed slightly reduced growth when cultivated with 1 µM Fe supplementation and wt-like biomass production under +Fe conditions. As previously reported for *A. fumigatus* strain ATCC46645 (11), the *ΔsidL* mutant strain exhibited wt-like TAFC but reduced FC production under both -Fe and 1 µM Fe conditions. In contrast, but in agreement with previously reported results (7), *ΔsidF* failed to produce TAFC under both-Fe and 1 µM Fe conditions. Interestingly, *ΔsidF* exhibited increased cellular FC accumulation, which was more pronounced under -Fe compared to 1 µM Fe conditions. These results indicate an interdependence between FusTS and FchTS production. The *ΔsidLΔsidF* strain failed to produce both TAFC and FC, similar to *ΔsidA.* The lack of siderophore production of *ΔsidLΔsidF* is in line with a *ΔsidA*-like growth pattern, i.e., the failure to grow under severe iron limitation. Taken together, growth and siderophore production results indicate that SidF is the previously unknown FchTS biosynthetic enzyme that complements SidL for FC production during iron limitation. Consistent with the previously predicted induction of the SidL complementing enzyme by iron limitation (11), SidF has been shown to be transcriptionally repressed by iron via SreA (7,28).

**Figure 3:**
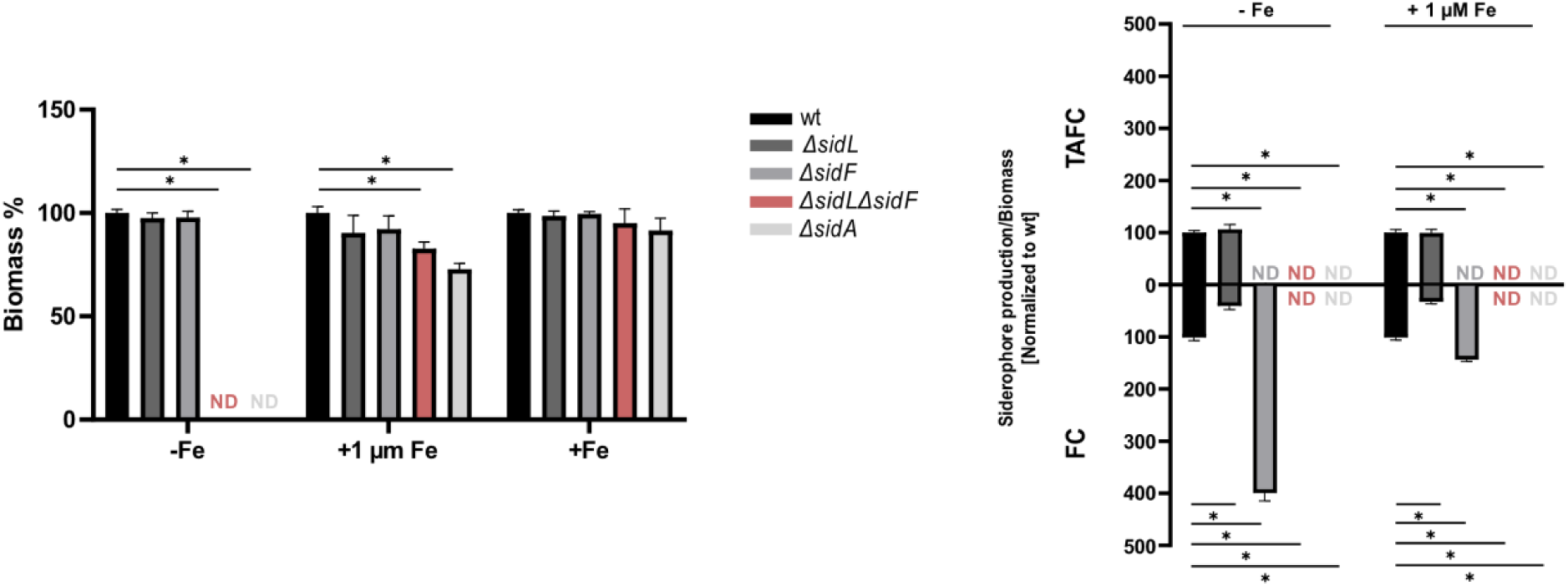
Lack of SidL decreases FC production, lack of SidF abolishes TAFC production and increases FC production, while lack of both enzymes abolishes production of both TAFC and FC, similar to lack of SidA. 100 mL shake flask cultures were inoculated with 1 × 10^8^ conidia of the respective fungal strain. The used minimal medium lacked iron addition (-Fe) or was supplemented with FeSO_4_ to a final concentration of 1 µM (+ 1 µM Fe) or 30 µM (+Fe). After incubation at 37 °C for 24 h, the biomass and supernatant of each culture was collected. The biomass was measured after freeze-drying and normalized to wt grown under the same condition. TAFC was extracted from collected supernatants and FC from freeze-dried biomasses from iron limited cultures (-Fe and 1 µM Fe); values were normalized to that of wt grown under the same condition. Siderophore production under iron limitation (-Fe and 1 µM Fe) was normalized first to biomass and then to wt. Under iron sufficiency, siderophore production was not detected for any strain. Absence of growth and siderophore detection is indicated by ND. The figure shows the means ± SD of biological triplicates; statistically significant differences (p ≤ 0.005) to wt calculated by one-way ANOVA are indicated with *. Absolute values are given in Supplementary Table S1.

### Both the N-and the C-terminal domains of SidF are essential for function in both FusTS and FchTS biosynthesis

SidF has been shown to consist of two structurally similar and independently folded domains, the N-terminal unique domain and the C-terminal GNAT motif-containing domain showing similarity to SidL, which are connected by a linker region (29). Consistent with the presence of the GNAT motif, the C-terminal domain, but not the N-terminal domain, has been shown to retain enzymatic activity *in vitro*.

To analyze the role of the two SidF domains for the enzymatic function, mutant strains expressing either full-length SidF, only the N-terminal SidF domain (amino acid residues 1-200; indicated in the strain term as *^Nterm(AKL^*^)^) or only the C-terminal domain (amino acid residues 201-462; indicated in the strain term as *^Cterm^*) under control of xylose-inducible P*xylP* promoter (indicated in the strain term as *^X^*) (31) with or without N-terminal tagging with the yellow fluorescence protein derivative Venus (indicated in the strain term as *^V^*) (32). Here, only N-terminal tagging could be applied as SidF localizes to peroxisomes via its C-terminal PTS1 motif. Cytosolic mislocalization has been shown to block SidF function in FusTS biosynthesis as long as SidI and SidH are localized to peroxisomes (20). The exclusive C-terminal SidF domain carries its natural PTS1 motif (AKL), while the exclusive N-terminal domain was artificially C-terminally tagged with the PTS1 motif AKL (indicated as *^(AKL)^*) for peroxisomal targeting. All three *sidF* alleles were expressed in a *ΔsidLΔsidF* genetic background, which allows analysis of SidF functionality by simple growth assays. Northern blot analysis confirmed the xylose-inducible expression of all *sidF* alleles at the transcriptional level, independent of iron availability (Supplementary Figure S1); Western blot analysis using a GFP-specific antibody confirmed the expression of all Venus-tagged *sidF* alleles at the protein level, independent of iron availability (Supplementary Figure S2). Notably, the C-terminal SidF domain showed significantly reduced protein levels compared to both full-length SidF and the N-terminal SidF domain, indicating reduced protein stability as all alleles were expressed under control of P*xylP* and showed similar transcript levels.

As expected, Venus-tagged full-length SidF (*ΔsidL,sidF^X,V^*), the N-terminal SidF domain (*ΔsidL,sidF^X,V,Nterm(AKL)^*) and the C-terminal SidF domain (*ΔsidL,sidF^X,V,Cterm^*) localized to punctate dots, which have previously been shown to be peroxisomes (20), demonstrating the intended localization of these SidF alleles, even of the allele carrying the artificial PTS1 sequence (Figure 4). Notably, the C-terminal SidF domain showed an about 5-fold lower fluorescence intensity compared to both full-length SidF and the N-terminal SidF domain, which is in line with the reduced protein stability observed in Western blot analysis (Supplementary Figure S2). In comparison, the generated strain *ΔsidF,sidL^X,V^*expressing N-terminal Venus-tagged SidL under P*xylP* control, showed cytosolic localization of SidL, as previously observed in the ATCC46645 background (11) .

**Figure 4:**
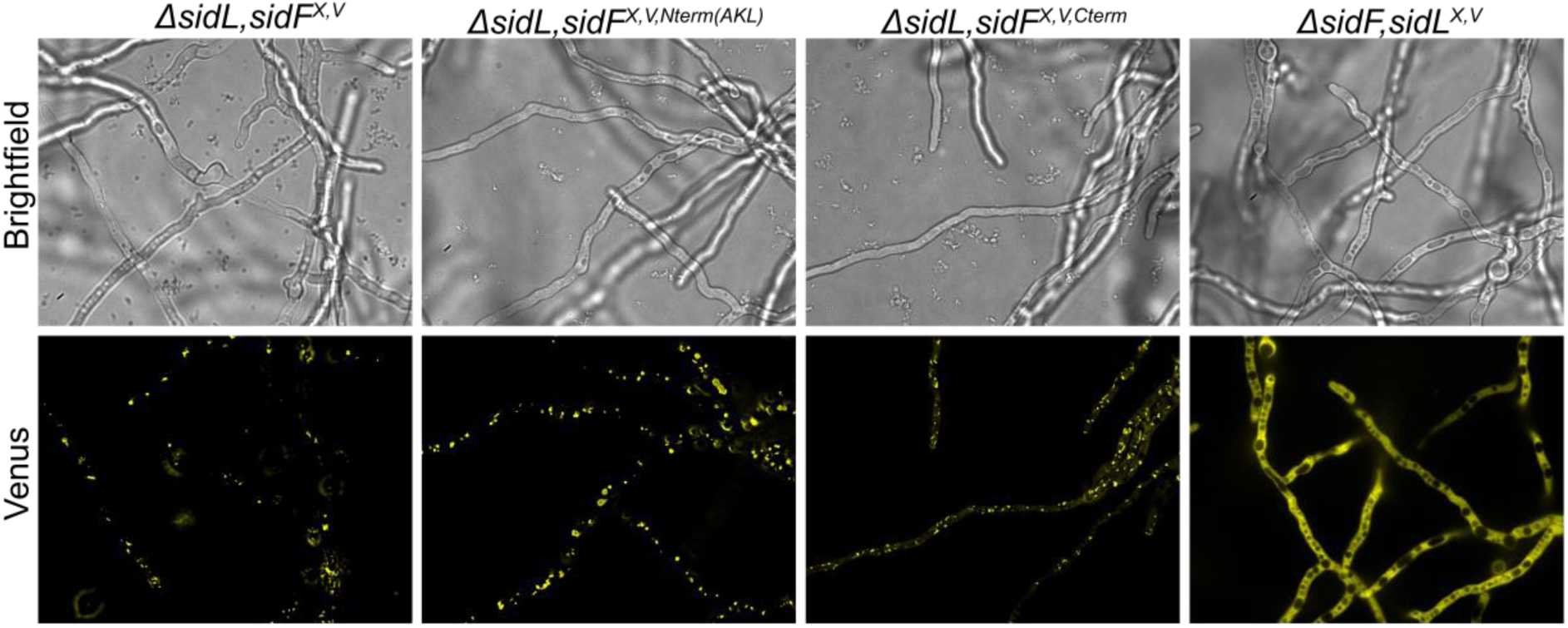
Venus-tagged full-length SidF (*ΔsidL,sidF^X,V^*), the N-terminal SidF domain (*ΔsidL,sidF^X,V,Nterm(AKL)^*) and the C-terminal SidF domain (*ΔsidL,sidF^X,V,Cterm^*), localize to peroxisomes, whereas Venus-tagged SidL localizes to the cytoplasm. Fungal strains were grown for 16 hours at 37 °C in 8-well chamber slides with 200 µl of minimal medium supplemented with 30 μm iron and 0.1 % xylose, using an inoculum of 10^4^ spores. Notably, the image intensity of strain *ΔsidL,sidF^X,V,Cterm^* was increased 5-fold to detect protein localization. Strain *ΔsidF,sidL^X,V^* is shown as example for a cytosolic localized protein, N-terminal Venus-tagged SidL.

In contrast to wt and *ΔsidL,sidF^X,V^* strains, *ΔsidL,sidF^X,V,Nterm(AKL)^* and *ΔsidL,sidF^X,V,Cterm^*showed an impaired growth on solid media under iron limitation (-Fe and 1 µM Fe + BPS) with and without xylose supplementation, phenocopying *ΔsidFΔsidL* (Supplementary Figure S3). These results strongly indicate that neither the N-nor the C-terminal domain of SidF (with N-terminal Venus tag), confers acyltransferase activity for the biosynthesis of FusTS or FchTS *in vivo*. The strains expressing the very same N-or C-terminal SidF domains but without the N-terminal Venus tag (*ΔsidL,sidF^X,Nterm(AKL)^* and *ΔsidL,sidF^X,Cterm^*) showed the same growth defect (Supplementary Figure S4), indicating that the N-terminal Venus tag does not negatively interfere with the enzymatic functionality.

Remarkably, the *ΔsidL,sidF^X,V^* strain showed similar biomass formation compared to wt and *ΔsidL* strains in liquid shake flask cultures without iron addition (-Fe), even without xylose supplementation (Figure 5A). In line with the growth pattern, this strain showed TAFC production under iron limitation (-Fe and 1 µM Fe), although lower than *ΔsidL.* Northern and Western blot analyses failed to detect expression of full-length SidF without xylose induction in *ΔsidL,sidF^X,V^*(see above). Therefore, these results demonstrate that already very low SidF amounts, due to the leakiness of the *xylP* promoter (31), are sufficient to rescue the growth defect and to enable limited TAFC biosynthesis. With xylose supplementation, *ΔsidL,sidF^X,V^*showed *ΔsidL*-like production of TAFC and FC (Figure 5B), proving full functionality of the P*xylP* controlled full-length *sidF* allele. The *ΔsidL,sidF^X^*strain, which expresses full-length *sidF* without the Venus tag, showed the same growth and siderophore production pattern as the *ΔsidL,sidF^X,V^* strain, which produces the Venus-tagged SidF (Supplementary Figure S5), corroborating the results. Similar to the growth on solid medium, *ΔsidL,sidF^X,V,Nterm(AKL)^* and *ΔsidL,sidF^X,V,Cterm^* showed a growth defect in liquid shake flask cultures without the addition of iron (-Fe) and irrespectively of xylose supplementation (Figure 5). Consistent with the growth pattern, these two strains lacked production of both TAFC and FC (1 µM Fe supplementation) even with xylose supplementation (Figure 5). Taken together, these results demonstrate that neither the N-nor the C-terminal domain of SidF, tagged with Venus, individually confer acyltransferase activity for biosynthesis of FusTS or FchTS *in vivo*. The *ΔsidL,sidF^X,Nterm(AKL)^* and *ΔsidL,sidF^X,Cterm^*strains, which express the individual N-and C-terminal SidF domains without Venus tag, displayed the very same growth and siderophore production pattern as the Venus-tagged counterparts (Supplementary Figure S5), reinforcing the results.

**Figure 5:**
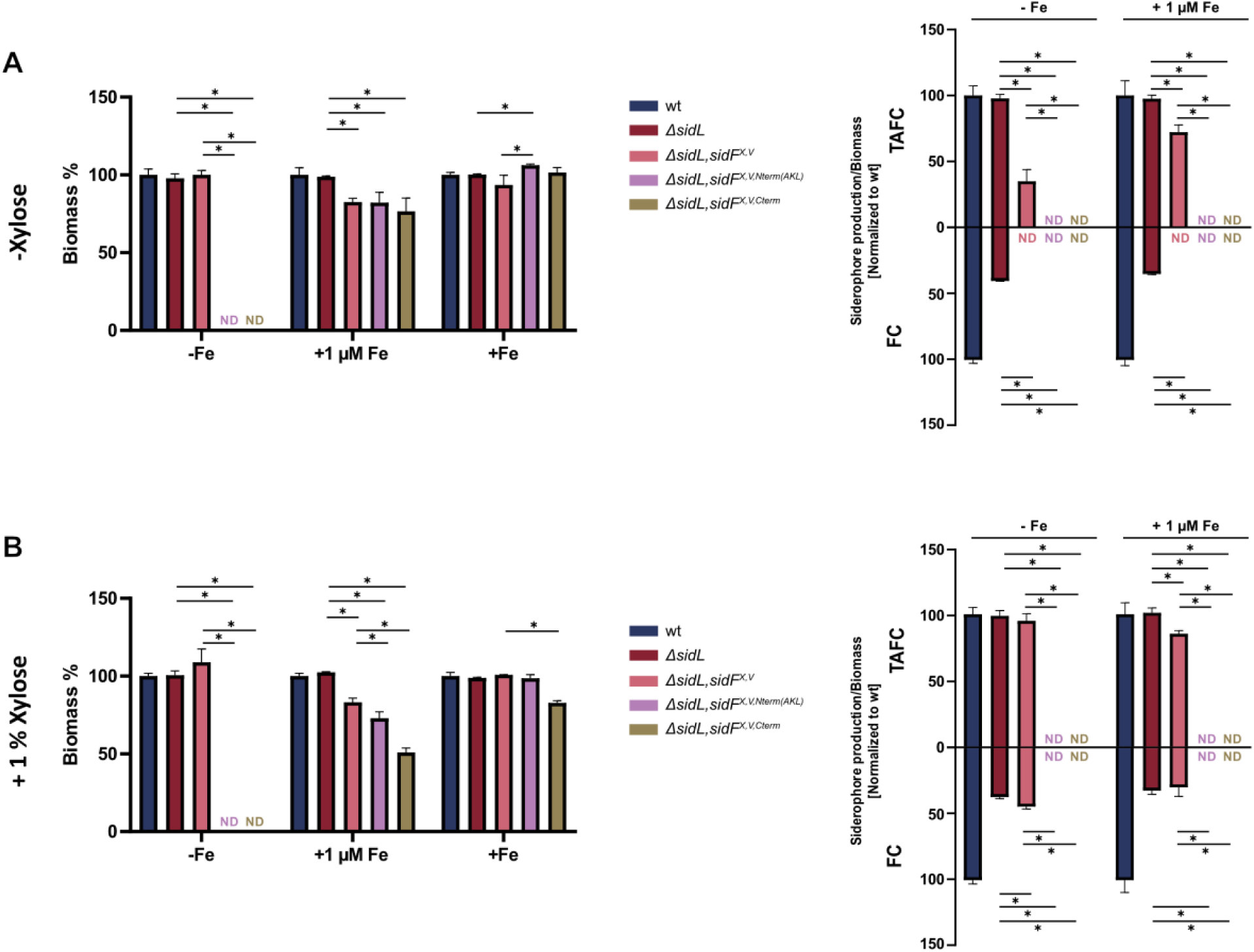
Neither the individual N-nor the C-terminal domain of SidF, tagged with Venus, supports biosynthesis of FusTS or FchTS *in vivo*. Liquid shake flask cultures and determination of siderophore production were done as described in Figure 3. However, cultures were either not supplemented (A) or supplemented with 1 % xylose for P*xylP* induction (B). Statistically significant differences were calculated by one-way ANOVA (p ≤ 0.005) compared to *ΔsidL* and *ΔsidL,sidF^X,V^*, respectively, and are indicated with *. Absolute values are given in Supplementary Table S2.

### PTS1 sequence tagging of SidL does not support FusST biosynthesis

As the data shown above indicates, that SidF is a bifunctional enzyme supporting the production of both FusTS and FchTS, the question arises as to whether SidL might also be bifunctional. In a previous study (7) and consistent with the data presented in Figure 3, it has been shown, that a mutant lacking SidF is unable to produce FusTS, i.e. that SidL is unable to support FusTS biosynthesis. However, as mentioned above, the peroxisomal localization of SidF has been shown to be essential for FusTS biosynthesis, as it must be present in the same compartment as SidH and SidI (20), whereas SidL localizes to the cytosol, as shown previously (11) and confirmed here (Figure 4). Consequently, peroxisomal localization of the acyltransferase may be essential for supporting FusTS biosynthesis. Therefore, we aimed to target the P*xylP*-controlled and N-terminal Venus-tagged SidL to the peroxisomes by artificial C-terminal tagging with a PTS1 motif (AKL) in a mutant lacking SidF. Western blot analysis demonstrated that P*xylP*-mediated regulation allows xylose-inducible expression of N-terminal Venus-tagged SidL without (*ΔsidF,sidL^X,V^*) and with PTS1 sequence tagging (*ΔsidF,sidL^X,V,(AKL)^*) (Supplementary Figure S6). However, *ΔsidF,sidL^X,V,(AKL)^* still showed predominant cytosolic localization of the PTS1-tagged SidL variant, similar to *ΔsidF,sidL^X,V^* (Supplementary Figure S7). As it cannot be excluded that fractions of SidL localize to peroxisomes, we still analyzed the consequences of the PTS1 tagging of SidL. In contrast to wt and *ΔsidF*, both *ΔsidF,sidL^X,V^*and *ΔsidF,sidL^X,V,(AKL)^* showed a growth defect on solid media under iron limitation (-Fe and 1 µM Fe + BPS) without xylose supplementation, similar to *ΔsidFΔsidL* (Supplementary Figure S8). Supplementation with xylose, partially cured the growth defect but could not restore *ΔsidF-* like growth, suggesting a decreased SidL function of the P*xylP*-driven and Venus-tagged SidL allele. Interestingly, *ΔsidF,sidL^X,V,(AKL)^* showed a more prominent growth defect than *ΔsidF,sidL^X,V^* with and without xylose, suggesting an impaired SidL function due to the PTS1 tagging. In liquid shake flask culture, the mutant expressing SidL under control of P*xylP* (*ΔsidF,sidL^X,V^*) was able to grow and to produce small amounts of FC without xylose supplementation under iron limitation (-Fe), most likely due to the aforementioned leakiness of the *xylP* promoter (Figure 6A). In contrast, *ΔsidF,sidL^X,V,(AKL)^*was unable to grow under this condition (Figure 6A), like *ΔsidLΔsidF* (see above), indicating that PTS1 sequence-tagging impairs SidL activity. Xylose supplementation increased FC production of *ΔsidF,sidL^X,V^*, but not to the levels produced by *ΔsidF* under -Fe conditions (Figure 6B), confirming that the P*xylP*-driven and Venus-tagged allele confers lower support for FchST biosynthesis. Xylose supplementation allowed growth of *ΔsidF,sidL^X,V,(AKL)^*, however to a lower degree compared to *ΔsidF* and *ΔsidF,sidL^X,V^*, which confirms, that PTS1-tagging impairs SidL function. Taken together, these studies showed, that C-terminal tagging of SidL with the PTS1 motif (AKL) does not compensate for the lack of *sidF* (*ΔsidF,sidL^X,V,(AKL)^*) and fails to support TAFC biosynthesis.

**Figure 6:**
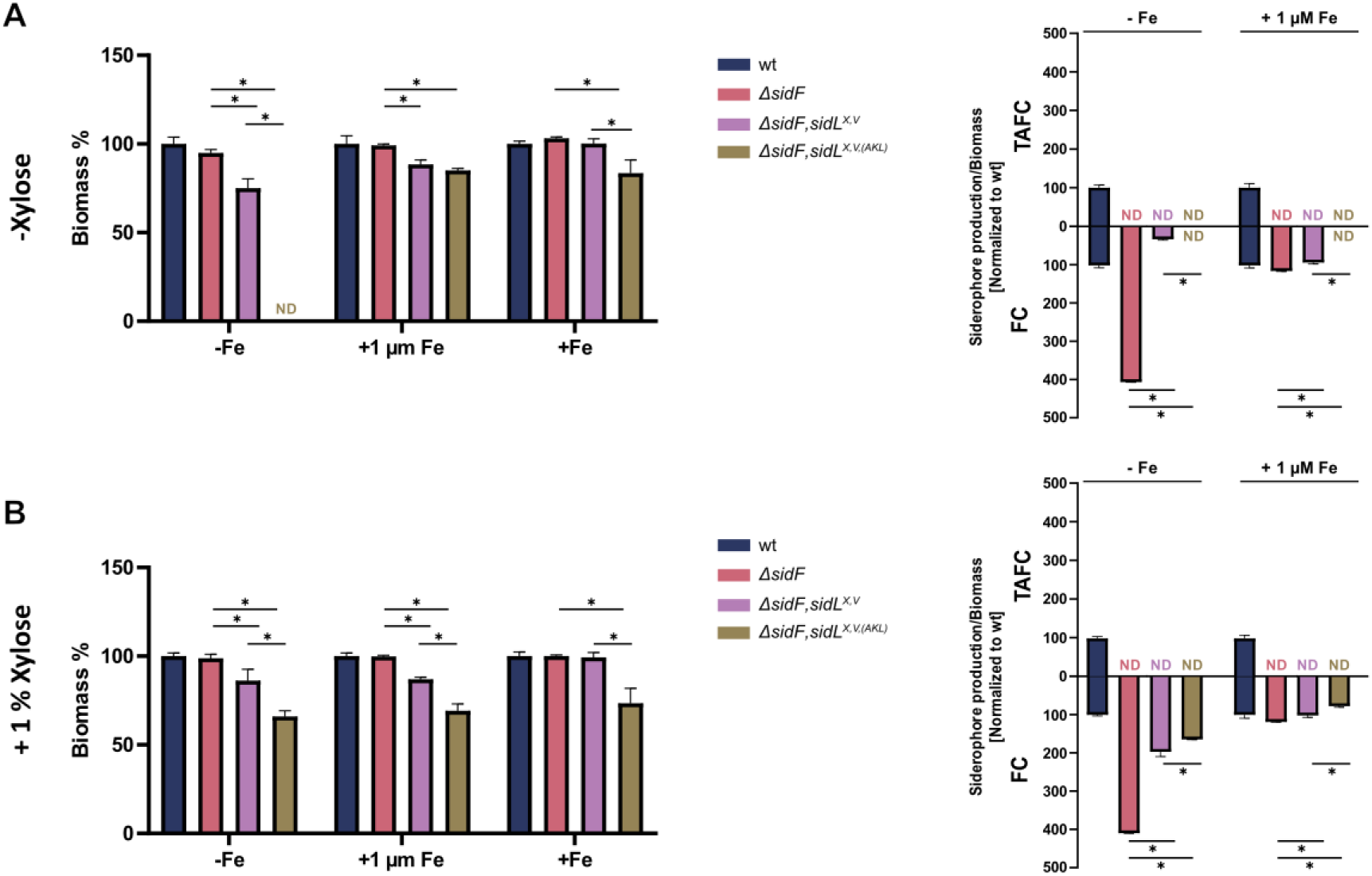
PTS1 sequence tagging impairs SidL activity and does not support FusST biosynthesis. Liquid shake flask cultures and determination of siderophore production were done as described in Figure 3. Cultures were either not supplemented (A) or supplemented with 1 % xylose for P*xylP* induction (B). Statistically significant differences were calculated by one-way ANOVA (p ≤ 0.005) compared to *ΔsidF* and *ΔsidF, sidL^X,V^*, respectively, and are indicated with *. Absolute values are given in Supplementary Table S4.

### SidL does not support FusTS biosynthesis, even in the absence of peroxisomes

Mislocalization of one or two of the three peroxisomal enzymes SidI, SidH and SidF to the cytosol has been demonstrated to impair FusTS biosynthesis in both *A. fumigatus* and *A. nidulans* (20). However, absence of peroxisomes, e.g., by inactivation of the peroxisome biogenesis protein PexC (*ΔpexC*), did not block FusTS biosynthesis in *A. nidulans* (20), indicating that FusTS biosynthesis requires the three enzymes SidI, SidH and SidF to localize to the same cellular compartment. As it was not possible to efficiently target SidL to peroxisomes by PTS1 motif tagging, we investigated whether SidL is able to support FusTS biosynthesis in the absence of peroxisomes. Therefore, we generated three *A. fumigatus* mutant strains lacking peroxisomes (*ΔpexC*), peroxisomes and SidF (*ΔpexCΔsidF*), and peroxisomes and SidL (*ΔpexCΔsidL*). In *A. fumigatus,* inactivation of PexC blocked ß-oxidation and consequently the utilization of fatty acids as carbon source, decreased conidial pigmentation and caused biotin auxotrophy (Supplementary Figure S9), as previously shown for *A. nidulans* (20,33–35). Reintegration of the *pexC* gene in the *ΔpexC* strain cured all defects (Supplementary Figure S10), demonstrating that the defects are specific to the *pexC* deletion. In liquid shake flask cultures, *ΔpexC*, *ΔpexCΔsidF* and *ΔpexCΔsidL* strains exhibited largely similar biomass formation compared to wt, *ΔsidF* and *ΔsidL* strains under iron limitation (-Fe and 1 µM Fe supplementation; Figure 7). Interestingly, PexC deficiency slightly increased biomass formation under iron sufficient conditions (30 µM Fe). Remarkably, the lack of peroxisomes (*ΔpexC*) significantly increased the production of FC, especially under severe iron limitation (-Fe), which was comparable to the FC production detected in the mutant strains lacking SidF (*ΔsidF*), PexC and SidF (*ΔpexCΔsidF*) and, to a lower degree PexC and SidL (*ΔpexCΔsidL*) (Figure 7). These data indicate that peroxisomal localization of the FusTS biosynthetic enzymes is important for optimization of TAFC biosynthesis. Furthermore, *ΔpexCΔsidF* did not produce TAFC, suggesting that SidL is unable to support FusTS biosynthesis, even in the absence of peroxisomes. In contrast, SidF was able to support biosynthesis of both FusTS and FchTS with both peroxisomal (*ΔsidL*) and cytosolic localization (*ΔpexCΔsidL*).

**Figure 7:**
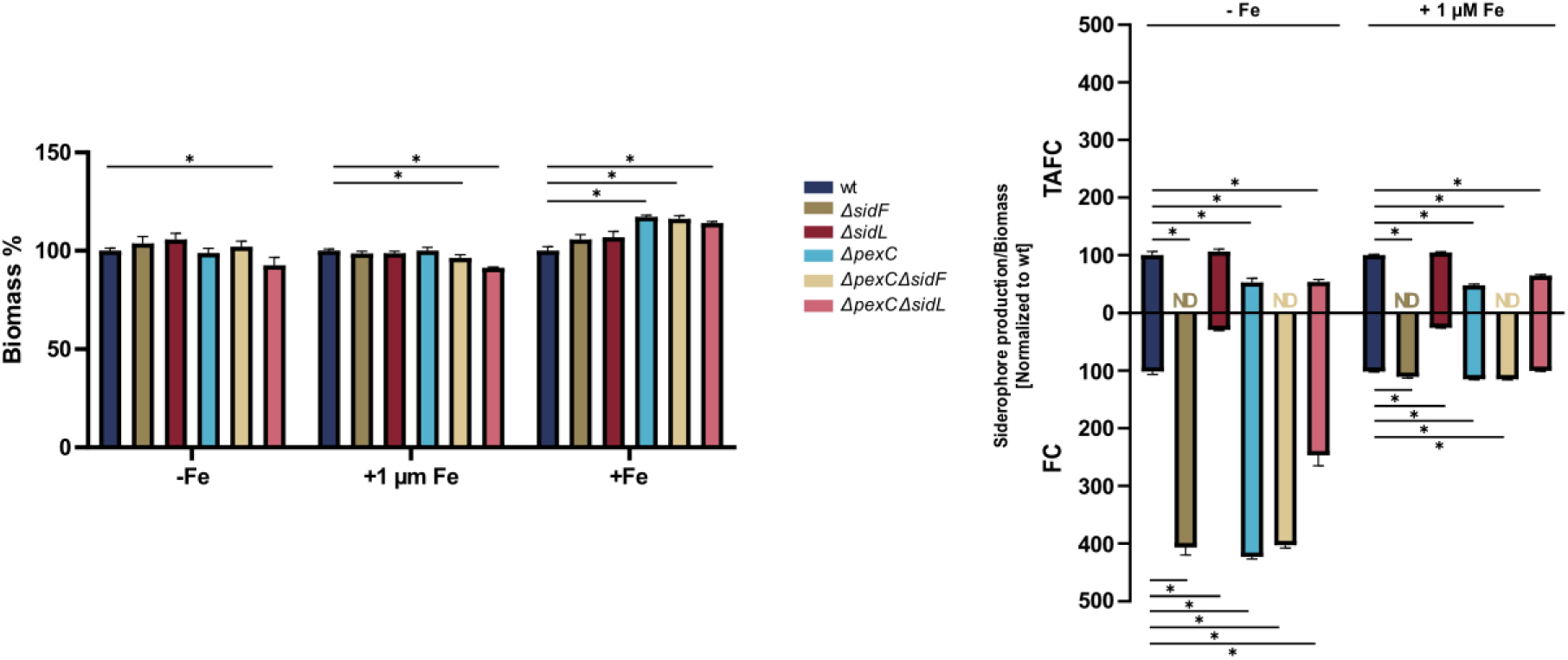
SidL does not support FusTS biosynthesis even in the absence of peroxisomes. Liquid shake flask cultures and determination of siderophore production were done as described in Figure 3. All cultures were supplemented with biotin due to biotin auxotrophy of *ΔpexC* strains (Supplementary Figure S9). Statistically significant differences were calculated by one-way ANOVA (p ≤ 0.005) compared to wt and are indicated with *. Absolute values are given in Supplementary Table S5.

### *A. fumigatus* SidF and SidL belong to two different subclades displaying differences in the GNAT consensus sequences

Blastp searches revealed homologs of SidF and/or SidL in 58 out of 82 included fungal species including many Ascomycetes and Basidiomycetes known to synthesize hydroxamate-class siderophores (Supplementary Figure S11), while no homologs were found in species that lack synthesis of hydroxamante-class siderophores such as Saccharomycotina, *Cryptococcus neoformans*, Mucoromycota, and Chytridiomycota. Therefore, the presence of SidF or SidL homologs appears to be indicative for the ability to produce siderophores similar to the presence of SidA (36). Phylogenetic analysis revealed that *A. fumigatus* SidF and SidL represent members of two phylogenetically separate clades, termed SidF and SidL clade, respectively (Supplementary Figure S11), suggesting an early divergence of the SidF and SidL clades. Interestingly, there are some outliers such as the N^5^-hydroxyornithine transacetylase Sib3 (NP_595895), which is involved in ferrichrome biosynthesis in *Schizosaccharomyces pombe* (37). Notably, phylogenetic analysis of the extracted GNAT domains of the SidF and SidL homologs revealed largely the same clustering as found for full-length SidF and SidL homologs (Supplementary Figure S12). In line with the clustering, SidF and SidL GNAT domains exhibit overlapping but distinct consensus sequences (Figure 8 and Supplementary Figure S13). These data may indicate the molecular determinants for their overlapping but distinct substrate specificity, acetyl-CoA for the SidL clade and both acetyl-CoA and anhydromevalonyl-CoA for the SidF clade. Noteworthy, the histidine residue previously shown to be essential for activity of siderophore biosynthetic N^6^-hydroxylysine acetyltransferase Rv1347c from *Mycobacterium tuberculosis* is conserved in both the SidF and the SidL clade acyltransferases (38). An alignment of the GNAT domains of *M. tuberculosis* Rv1347c with *A. fumigatus* SidF and SidL is shown in (Supplementary Figure S14).

**Figure 8.**
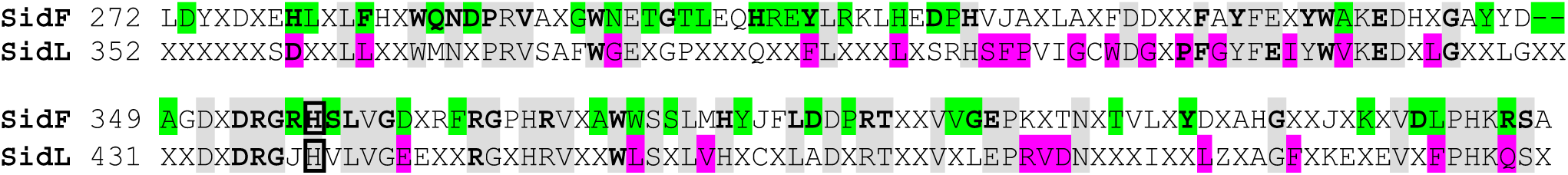
SidF and SidL clade GNAT domains reveal distinct consensus sequences. Numbering is based on *A. fumigatus* SidF and SidL. The consensus sequence logos of SidF and SidL clades are shown in Supplementary Figure SX3 and were extracted from the multiple alignment of the phylogenetic tree shown in Supplementary Figure SX2 using Geneious Prime. Shown are amino acid residues with an identity of >50% within the respective clade. Amino acid residues conserved in both clades are highlighted in grey; amino acid residues with >80% unique conservation within their clades are highlighted in green for the SidF clade and in pink for the SidL clade. Amino acid residues with >99% conservation are shown in bold. The histidine residue shown to be essential for activity of siderophore biosynthetic N^6^-hydroxylysine acetyltransferase Rv1347c from *Mycobacterium tuberculosis* is boxed (38).

## DISCUSSION

Most fungal species employ SIA, with the most commonly used hydroxamate-class siderophores being FusTS, FchTS and coprogen-type siderophores (3,6). After the first and common step in the biosynthesis of hydroxamate-class siderophores, SidA-catalyzed formation of N^5^-hydroxyornithine (5), the pathway diverges depending on the siderophore type. Biosynthesis of FusTS and coprogen-type siderophores requires the incorporation of an anhydromevalonyl group to form N^5^-anhydromevalonyl-N^5^-hydroxyornithine, which in *A. fumigatus* is exclusively catalyzed by the transacylase SidF (7). For the biosynthesis of FchTS, N^5^-hydroxyornithine is acetylated to N^5^-acetyl-N^5^-hydroxyornithine, in *A. fumigatus* mediated by the constitutively active transacetylase SidL and by a previously unidentified enzyme that complements this function during iron starvation (11). Structural characterization of SidF revealed a two-domain architecture with tetrameric assembly and the comparison with the AlphaFold model of SidL indicated structural similarity between SidF and SidL (29). SidF and SidL share only limited amino acid sequence similarity in their C-terminal GNAT (Gcn5-related N-acetyltransferases) motif-containing domains, but not in their N-terminal domains. Furthermore, SidF and SidL show other significant differences (11): SidF is transcriptionally induced by iron limitation and localizes to peroxisomes (7,20,28), whereas SidL is a cytosolic enzyme with largely iron-independent expression (11).

Here we show that the simultaneous absence of both SidF and SidL blocks biosynthesis of both the FusTS TAFC and the FchTS FC and that the respective mutant phenocopies the mutant lacking SidA, i.e., it shows a severe growth defect during iron limitation. These results strongly suggest that SidF is a bifunctional enzyme accepting not only anhydromevalonyl-CoA but also acetyl-CoA as substrates for the biosynthesis of FusTS and FchTS (Figure 1). In line with this, *in vitro* biochemical studies showed that a recombinant SidF protein accepts acetyl-CoA as a substrate, with the catalytic activity mediated by the C-terminal domain containing the GCN5-related N-acetyltransferase (GNAT) motif (29). However, using independent expression of the individual N-and C-terminal SidF domains in peroxisomes of *A. fumigatus,* we show that *in vivo* biosynthesis of both FusTS and FchTS depends on a full-length SidF protein containing both domains. In this context, *in vitro* studies have shown that the N-terminal SidF domain contributes to protein solubility and oligomerization, while the C-terminus mediates enzymatic activity and oligomer formation (29). Furthermore, it cannot be excluded that the N-terminal SidF domain mediates interaction with other proteins required for siderophore biosynthesis. Interestingly, expression in *A. fumigatus* indicated that the individual C-terminal SidF domain has a significantly reduced protein stability compared to its N-terminal domain.

Since SidF and SidL are structurally similar and share the GNAT motif, we tested whether SidL might be a dual substrate enzyme like SidF, supporting the biosynthesis of both FusTS and FchTS. A previous study (7) and the data presented here clearly show that a mutant lacking SidF is unable to produce FusTS, i.e., that SidL is unable to support FusTS biosynthesis. However, peroxisomal localization of SidF has been shown to be essential for FusTS biosynthesis, as it must be present in the same compartment as SidH and SidI (20), while SidL shows cytosolic localization as shown previously (11) and confirmed here. Since the peroxisomal localization of the acyltransferase SidF is essential for supporting FusTS biosynthesis, we tried to target SidL to peroxisomes by artificial C-terminal tagging with a PTS1 (AKL) sequence. However, this approach failed to efficiently localize SidL to peroxisomes and the engineered SidL did not support FusTS biosynthesis in the corresponding strains.

Although mislocalization of SidF to the cytosol has been shown to impair FusTS biosynthesis in both *A. fumigatus* and *A. nidulans*, lack of peroxisomes, did not block FusTS biosynthesis in *A. nidulans* (20), suggesting that FusTS biosynthesis requires SidF to colocalize with the other two originally peroxisomal FusTS enzymes, SidI and SidH, in the same cellular compartment. However, even in strains lacking peroxisomes due to inactivation of the peroxin PexC, SidL did not support FusTS biosynthesis, strongly suggesting that SidL does not accept anhydromevalonyl-CoA as a substrate for acylation of N^5^-hydroxyornithine. The fact that SidF and SidL are structurally similar, share the GNAT motif, and have overlapping but not identical substrate specificities may help to elucidate the molecular determinants for their substrate specificity.

Lack of peroxisomes reduced the production of the FusTS TAFC, concomitant with increased production of the FchTS FC. In other words, peroxisomal localization of FusTS enzymes is important for optimizing FusTS production at the expense of FchTS. It is tempting to hypothesize that the latter is caused by the lack of compartmentalization of the peroxisomal FusTS-specific enzymes with the FusTS precursor mevalonate (Figure 1).

Lack of SidF blocked the biosynthesis of TAFC but significantly increased the production of FC, suggesting interdependence between FusTS and FchTS production. In wt, production of TAFC exceeds that of FC about 10-fold (Supplementary Tables S1 -S5). Therefore, this effect is likely to be caused by a change in precursor availability as the FusTS precursor mevalonate is derived from acetyl-CoA and the blocked FusTS biosynthesis might increase acetyl-CoA availability for biosynthesis of FchTS.

In fungi, SidF and SidL homologs are found exclusively in species capable of synthesizing hydroxamate-class siderophores. SidF and SidL homologs, as well as extracted GNAT domains, group into phylogenetically independent clades. Their GNAT domains have overlapping but distinct consensus sequences which may reflect the molecular determinants for their overlapping but distinct substrate specificity, acetyl-CoA for the SidL clade and both acetyl-CoA and anhydromevalonyl-CoA for the SidF clade. SidF homologs are not only essential for biosynthesis of FusTS but also for coprogen-type siderophores, which also require incorporation of anhydromevalonyl moieties. Therefore, the findings of this study are likely to be relevant not only for other FusTS producing but also for coprogen-type siderophore producing fungal species.

Absence of SidA has been shown to cause avirulence in several infection models including murine aspergillosis models (3,5,21,22,39,40). Therefore, SidA, and consequently siderophore biosynthesis, is a promising target for development of novel antifungal drugs. Here we demonstrated that simultaneous inactivation of SidF and SidL phenocopies the inactivation of SidA. SidL and SidF share similarity in the C-terminal GNAT motif containing domain and the common substrate acetyl-CoA. Therefore, simultaneous targeting of SidF and SidL is an alternative to SidA inhibition for development of novel antifungal drugs, with the same outcome – inhibition of biosynthesis of all siderophore types – but targeting a different class of enzymes. In conclusion, this study has uncovered an additional role for SidF in fungal siderophore biosynthesis, which may be instrumental for the identification of molecular determinants of substrate specificity of GNAT enzymes, revealed the interdependence of production of FusTS and FchTS production and shed light on the rational of peroxisomal localization of enzymes required for FusTS biosynthesis. In addition, the results reinforced the potential of the siderophore biosynthetic enzymes SidF and SidL as targets for the development of new antifungal therapies.

## MATERIAL AND METHODS

### Growth Conditions

For spore production, *Aspergillus fumigatus* strains were grown in *Aspergillus* complex media containing 1 g/L yeast extract (Lab M Limited, Rochdale, UK), 20 mg/L Glucose (Glc) (Carl Roth, Karlsruhe, Germany), 1 g/L casamino acids (Sigma-Aldrich Inc., St. Louis, MO, USA), 2 g/L peptone (Carl Roth GmbH, Karlsruhe, Germany), salt solution, and iron-free trace elements (41), at 37°C for 5 days. For all other experiments, *Aspergillus* minimal medium containing 1% (*w/v*) glucose as carbon source and 20 mM glutamine (Gln) as nitrogen source was used (41); the trace element solution was prepared without iron salts; FeSO_4_ supplementation was performed as described in the respective figures. For induction of the *xylP* promoter, xylose was added to a final concentration of either 0.1% or 1% (*w/v*). For plate growth essays, 10^4^ spores of each strain were point-inoculated onto plates and incubated for 48 h at 37 °C. 100 mL liquid shake flask cultures were inoculated with 10^6^ spores per mL and incubated with 200 rpm rotation at 37 °C for time periods described in the respective figures.

#### Generation of A. fumigatus strains

All strains used in this study (listed in Supplementary Table S6) were generated in *Aspergillus fumigatus* strain A1160+ (termed wt here), a derivative from the clinical isolate *A. fumigatus* CEA10, but lacking non-homologous recombination (ΔakuB^ku80^::*pyrG^−^zeo*, pyrG^−^::pyrG^Af^; MAT1-1) to facilitate genetic manipulation (30,42,43). All oligonucleotides used in this are summarized in Supplementary Table S7.

The SidF lacking mutant (*ΔsidF*) was generated by replacement of the *sidF* coding region by the hygromycin resistance cassette (*hph*) via homologous recombination according to (42). Therefore, a fragment comprising the 5′-NCR (non-coding region) of *sidF*, the hygromycin resistance cassette (*hph*) and the 3′-NCR of *sidF* was amplified by PCR from the genomic DNA (gDNA) of a previously published *ΔsidF* mutant strain (7). The SidL lacking mutant (*ΔsidL*) was generated by replacement of the *sidL* coding region by the pyrithiamine resistance cassette (*ptrA*). Therefore, the 5′-NCR of *sidL*, *ptrA* and the 3′-NCR of *sidL* were amplified by PCR, using gDNA as a template for the amplification of the NCRs and plasmid pSK275 for amplification of *ptrA* (44). Subsequently, the three fragments were linked by fusion PCR using nested primers. The mutant lacking both SidF and SidL (*ΔsidLΔsidF*) was generated by deletion of *sidL* as described above, but in the *ΔsidF* mutant strain.

The PexC lacking mutant (*ΔpexC*) was generated by replacement of the *pexC* coding region by *ptrA*. Therefore, the 5’-and 3’-NCRs of *pexC* and *ptrA* were amplified by PCR, using gDNA as template for amplification of the NCRs or plasmid pSK275 for *ptrA* (44). Subsequently, the three fragments were then linked by fusion PCR using nested primers. The mutant lacking both PexC and SidF (*ΔpexCΔsidF*) was generated by deletion of of *sidF* as described above but in the *ΔpexC* background. The mutant lacking both PexC and SidL was generated by replacement of *sidL* in *ΔpexC* using a fusion PCR-generated construct containing the 5′-NCR of *sidL*, *hph* and the 3′-NCR of *sidL*. Here, plasmid pAN7-1 was used as template for amplification of *hph* (45).

To generate mutant strains *ΔsidL,sidF^X,V^*, *ΔsidL,sidF^X,V,Nterm(AKL)^* and *ΔsidL,sidF^X,V,Cterm^*plasmids were generated that were integrated in *A. fumigatus ΔsidLΔsidF* at the *fcyB* locus, which allows positive selection with 5-flucytosine without the need for a selection marker gene (46). Therefore, four DNA fragments were amplified: (i) the plasmid backbone including 5°-and 3°-NCR of *fcyB* amplified from template p*fcyB* (46), (ii) P*xylP*-driven *venus*, amplified from template pMMHL69 (15), (iii) the 1389bp (full-length SidF), 600bp (N-terminal SidF domain including C-terminal PTS1 sequence), or 789 bp (C-terminal SidF domain) *sidF* fragments including 1000 bp of the *sidF* 3′-NCR, amplified from wt gDNA. To generate mutant strains *ΔsidF,sidL^X,V^* and *ΔsidF,sidL^X,V,(AKL)^* plasmids were generated that were integrated in *A. fumigatus ΔsidLΔsidF* at the *fcyB* locus (46). Therefore, four DNA fragments were amplified: (i) the plasmid backbone as described above (46), (ii) P*xylP*-driven *venus*, amplified from template pMMHL69 (15), (iii) the *sidL* coding gene including 1000 bp of its 3′-NCR or the *sidL* coding gene C-terminally tagged with PTS1 sequence and 1000 bp of the 3′-NCR, amplified from wt gDNA. To generate the mutant strain *pexC^Comp^* a plasmid was generated that was integrated in *A. fumigatus ΔpexC* at the *fcyB* locus (46). Therefore, two DNA fragments were amplified: (i) the plasmid backbone as described above (46), (ii) the *pexC* coding sequence including 960 bp of the 5’-NCR for amplification of the native promoter and 915 bp of the 3’-NCR, amplified from wt gDNA. The generated fragments were assembled using NEBuilder HiFi DNA assembly (New England Biolabs, Ipswich, MA, USA). Before transformation, the plasmids were linearized by restriction digestion using *Not*I.

PCR amplified transformation cassettes were purified by column purification (Monarch PCR and DNA Cleanup Kit, New England Biolabs, Ipswich, MA, USA) prior the use for fungal transformation of *A. fumigatus*. The transformation of *A. fumigatus* A1160+ was performed according to (47). Selection of transformants was carried out on AMM plates supplemented with 0.2 mg/mL hygromycin B, 0.1 μg/mL pyrithiamine or 10 μg/mL 5-flucytosine. Correct genetic manipulations were proven by Southern blot analysis (Supplementary Figures S15-S20) and growth assays.

### RNA Isolation and Northern Blot Analysis

RNA was isolated from mycelium using TRI reagent (Sigma-Aldrich Inc., St. Louis, MO, USA) as stated in the manufacturer’s protocol. 10 µg of total RNA were separated on a 1.2% agarose gel containing 1.85% (w/v) formaldehyde. Gels were blotted onto a Hybond^TM^-N membrane (Amersham Biosciences, Slough, UK). Detection by hybridization was performed using digoxigenin-labelled probes (PCR DIG Labelling Mix. Roche, Basel, Switzerland) amplified by PCR. The primers used for generation of the Northern blot hybridization probes are listed in Supplementary Table S8.

### Protein Isolation and Western Blot Analysis

Proteins were isolated by alkaline lysis from freeze-dried mycelium, followed by protein precipitation using trichloroacetic acid (48). 5 µl of total protein were separated on 10% SDS-PA gels which were either blotted onto a nitrocellulose membrane (Amersham^TM^ Protran ^TM^ Premium 0.45 µm NC, GE Healthcare) using a Trans-Blot turbo transfer system (Bio-Rad, Hercules, CA, USA) or used for Coomassie staining. Proteins of interest were detected using mouse α-GFP antibody (1:10,000 diluted; Roche, Basel, Switzerland, REF 11 814 460 001) as primary antibody in combination with peroxidase-coupled secondary anti-Mouse antibody (1:10,000 diluted; Sigma-Aldrich, St. Louis, MO, USA, A9044). ECL reagent (Amersham Biosciences, Slough, UK) was used for detection.

### Siderophore Analysis

Extraction and measurement of extracellular TAFC and intracellular FC was performed as previously described (24).

### Microscopy Analysis

The fungal strains were grown for 16 hours at 37 °C in 8-well chamber slides (µ-Slide 8 Well; Ibidi, Gräfelfing, Germany) with a total volume of 200 µl minimal medium supplemented with 30 µM FeSO_4_ and 0.1 % xylose (w/v), inoculated with 10^4^ spores per well. Images were obtained as described in (49), using a 60× TIRF objective (Plan-APOCHROMAT 60×/1.49 Oil, Nikon, Tokyo, Japan) mounted on an inverted microscope (Eclipse Ti2-E; Nikon, Tokyo, Japan) with a spinning disc confocal unit (CSU-W1, Yokogawa, Tokyo, Japan) and an EMCCD camera (iXon Ultra 888, Andor, Belfast, Ireland) with an additional 1.5× magnification, using the NIS-Elements software (Nikon, Tokyo, Japan). The Lucy–Richardson algorithm was applied for deconvolution and Fiji ImageJ (50) for processing of microscopy pictures.

### Bioinformatics

Proteins were blasted using Geneious Prime® 2024.0.7 (https://www.geneious.com), which accesses the NCBI database (51). Alignments were performed with Clustal Omega and phylogenetic trees were constructed using the Neighbor-Joining method using Geneious Prime.

### Statistical Analysis

Statistical analyses were performed with GrahPad Prism version 10.2.3 for Windows, GraphPad Software, San Diego, CA, USA, www.graphpad.com.

## ABBREVIATIONS

FC: ferricrocin
FchTS: ferrichrome-type siderophores
-Fe: iron limitation
+Fe: iron sufficiency
FusTS: fusarinine-type siderophores
gDNA: genomic DNA
NCR: non-coding region
NRPS: non-ribosomal peptide synthetase
PTS: peroxisomal targeting sequence
RIA: reductive iron assimilation
SIA: siderophore-mediated iron acquisition
TAFC: triacetylfusarinine C
wt: wild type

## Acknowledgements/funding

This research was funded by the Austrian Science Fund (FWF) [Grant-DOI:10.55776/W1253]. For open access purposes, the author has applied a CC BY public copyright license to any author accepted manuscript version arising from this submission. The authors acknowledge support from the Euregio Science Fund (SupErA IPN95 to HH and SB) and the Medical University of Innsbruck.

## Author contributions

P.C.: conceptualization, methodology, investigation, visualization, writing – original draft, writing – review & editing; A.Y.: conceptualization, methodology, supervision; S.O.: methodology, investigation; I.H.: methodology, investigation, visualization; T.P.: methodology, investigation; S. B.: conceptualization, funding, acquisition, project administration, supervision, writing – review & editing; H.H.: conceptualization, funding acquisition, project administration, supervision, visualization, writing – original draft, writing – review & editing, resources.

All authors have read and approved the final manuscript.

