## Supplementary Material for "*Aspergillus fumigatus* SidF is a dual substrate acyltransferase involved in biosynthesis of both fusarinine- and ferrichrome-type siderophores"

**Supplementary Table S1:** Absolute values (mean ± SD) of experiments presented in Figure 3 The TAFC content of the supernatant and the FC content of the mycelia were normalized to mg biomass. ND, not detected.

| Strain | Biomass (g) -Fe | Biomass (g) +1 µm Fe | Biomass (g) +Fe | TAFC (nmol/mg) -Fe | TAFC (nmol/mg) +1 µm Fe | FC (nmol/mg) -Fe | FC (nmol/mg) +1 µm Fe |
|---|---|---|---|---|---|---|---|
| **wt** | 0,125 ± 0,002 | 0,533 ± 0,016 | 0,678 ± 0,011 | 50,38 ± 2,18 | 18,81 ± 1,08 | 3,61 ± 0,21 | 2,6 ± 0,13 |
| ***ΔsidL*** | 0,122 ± 0,003 | 0,482 ± 0,045 | 0,669 ± 0,015 | 53,57 ± 4,55 | 18,65 ± 1,32 | 1,41 ± 0,25 | 0,8 ± 0,12 |
| ***ΔsidF*** | 0,122 ± 0,004 | 0,491 ± 0,034 | 0,675 ± 0,008 | ND | ND | 14,45 ± 0,54 | 3,8 ± 0,08 |
| ***ΔsidA*** | ND | 0,388 ± 0,015 | 0,621 ± 0,040 | ND | ND | ND | ND |
| ***ΔsidLΔsidF*** | ND | 0,441 ± 0,017 | 0,645 ± 0,047 | ND | ND | ND | ND |

**Supplementary Table S2:** Absolute values (mean ± SD) of experiments presented in Figure 5. The TAFC content of the supernatant and the FC content of the mycelia were normalized to mg biomass. ND, not detected.

| | Strain | Biomass (g) -Fe | Biomass (g) +1 µm Fe | Biomass (g) +Fe | TAFC (nmol/mg) -Fe | TAFC (nmol/mg) +1 µm Fe | FC (nmol/mg) -Fe | FC (nmol/mg) +1 µm Fe |
|---|---|---|---|---|---|---|---|---|
| **-Xylose** | **wt** | 0,104 ± 0,004 | 0,547 ± 0,025 | 0,659 ± 0,003 | 39,08 ± 2,87 | 10,86 ± 1,21 | 4,32 ± 0,12 | 2,87 ± 0,13 |
| | ***ΔsidL,sidF***$^{X,V}$ | 0,104 ± 0,003 | 0,451 ± 0,013 | 0,616 ± 0,041 | 13,65 ± 3,47 | 7,84 ± 0,60 | ND | ND |
| | ***ΔsidL,sidF***$^{X,V,Nterm(AKL)}$ | ND | 0,449 ± 0,036 | 0,700 ± 0,005 | ND | ND | ND | ND |
| | ***ΔsidL,sidF***$^{X,V,Cterm}$ | ND | 0,419 ± 0,046 | 0,670 ± 0,021 | ND | ND | ND | ND |
| | ***ΔsidL*** | 0,102 ± 0,002 | 0,540 ± 0,003 | 0,660 ± 0,004 | 38,2 ± 1,20 | 10,6 ± 0,30 | 1,73 ± 0,01 | 1,01 ± 0,01 |
| **+ 1 % Xylose** | **wt** | 0,111 ± 0,002 | 0,664 ± 0,012 | 0,678 ± 0,015 | 32,59 ± 1,68 | 10,86 ± 0,95 | 3,99 ± 0,12 | 3,45 ± 0,33 |
| | ***ΔsidL,sidF***$^{X,V}$ | 0,121 ± 0,009 | 0,552 ± 0,017 | 0,684 ± 0,001 | 30,98 ± 1,77 | 9,27 ± 0,27 | 1,77 ± 0,07 | 1,02 ± 0,24 |
| | ***ΔsidL,sidF***$^{X,V,Nterm(AKL)}$ | ND | 0,484 ± 0,027 | 0,669 ± 0,016 | ND | ND | ND | ND |
| | ***ΔsidL,sidF***$^{X,V,Cterm}$ | ND | 0,339 ± 0,018 | 0,562 ± 0,009 | ND | ND | ND | ND |
| | ***ΔsidL*** | 0,112 ± 0,003 | 0,68 ± 0,013 | 0,670 ± 0,002 | 32,2 ± 1,32 | 10,99 ± 0,40 | 1.51 ± 0,10 | 1,13 ± 0,10 |

**Supplementary Table S3:** Absolute values (mean ± SD) of experiments presented in Figure S5. The TAFC content of the supernatant and the FC content of the mycelia were normalized to mg biomass. ND, not detected.

| | Strain | Biomass (g) -Fe | Biomass (g) +1 µm Fe | Biomass (g) +Fe | TAFC (nmol/mg) -Fe | TAFC (nmol/mg) +1 µm Fe | FC (nmol/mg) -Fe | FC (nmol/mg) +1 µm Fe |
|---|---|---|---|---|---|---|---|---|
| **-Xylose** | **wt** | 0,144 ± 0,001 | 0,602 ± 0,003 | 0,670 ± 0,014 | 37,69 ± 2,39 | 8,97 ± 0,38 | 4,49 ± 0,02 | 2,85 ± 0,18 |
| | ***ΔsidL,sidF***$^{X,V}$ | 0,146 ± 0,001 | 0,529 ± 0,013 | 0,670 ± 0,003 | 17,22 ± 0,67 | 4,69 ± 0,11 | ND | ND |
| | ***ΔsidL,sidF***$^{X,V,Nterm(AKL)}$ | ND | 0,491 ± 0,018 | 0,632 ± 0,009 | ND | ND | ND | ND |
| | ***ΔsidL,sidF***$^{X,V,Cterm}$ | ND | 0,414 ± 0,004 | 0,635 ± 0,002 | ND | ND | ND | ND |
| | ***ΔsidL*** | 0,143 ± 0,004 | 0,588 ± 0,001 | 0,675 ± 0,003 | 36,79 ± 0,55 | 8,81 ± 0,02 | 1,83 ± 0,02 | 1,02 ± 0,05 |
| **+ 1 % Xylose** | **wt** | 0,158 ± 0,002 | 0,664 ± 0,008 | 0,722 ± 0,005 | 25,57 ± 1,48 | 5,30 ± 0,21 | 4,01 ± 0,05 | 3,65 ± 0,55 |
| | ***ΔsidL,sidF***$^{X,V}$ | 0,160 ± 0,002 | 0,625 ± 0,005 | 0,687 ± 0,027 | 23,31 ± 1,29 | 5,38 ± 0,28 | 1,75 ± 0,03 | 1,32 ± 0,05 |
| | ***ΔsidL,sidF***$^{X,V,Nterm(AKL)}$ | ND | 0,538 ± 0,015 | 0,647 ± 0,014 | ND | ND | ND | ND |
| | ***ΔsidL,sidF***$^{X,V,Cterm}$ | ND | 0,524 ± 0,009 | 0,639 ± 0,009 | ND | ND | ND | ND |
| | ***ΔsidL*** | 0,160 ± 0,001 | 0,691 ± 0,016 | 0,710 ± 0,017 | 24,85 ± 1,73 | 5,45 ± 0,49 | 1,51 ± 0,08 | 1,15 ± 0,09 |

**Supplementary Table S4:** Absolute values of experiments presented in Figure 6. The TAFC content of the supernatant and the FC content of the mycelia were normalized to mg biomass. ND, not detected.

| | Strain | Biomass (g) -Fe | Biomass (g) +1 µm Fe | Biomass (g) +Fe | TAFC (nmol/mg) -Fe | TAFC (nmol/mg) +1 µm Fe | FC (nmol/mg) -Fe | FC (nmol/mg) +1 µm Fe |
|---|---|---|---|---|---|---|---|---|
| **-Xylose** | **wt** | 0,104 ± 0,004 | 0,547 ± 0,025 | 0,659 ± 0,003 | 39,08 ± 2,87 | 10,86 ± 1,21 | 4,32 ± 0,12 | 2,87 ± 0,13 |
| | ***ΔsidF,sidL***$^{X,V}$ | 0,078 ± 0,006 | 0,484 ± 0,014 | 0,660 ± 0,018 | ND | ND | 1,84 ± 0,01 | 3,01 ± 0,06 |
| | ***ΔsidF,sidL***$^{X,V,(AKL)}$ | ND | 0,465 ± 0,006 | 0,551 ± 0,049 | ND | ND | ND | ND |
| | ***ΔsidF*** | 0,099 ± 0,002 | 0,542 ± 0,004 | 0,680 ± 0,005 | ND | ND | 17,50 ± 0,03 | 3,30 ± 0,03 |
| **+ 1% Xylose** | **wt** | 0,111 ± 0,002 | 0,664 ± 0,012 | 0,678 ± 0,015 | 32,59 ± 1,68 | 10,86 ± 0,95 | 3,99 ± 0,12 | 3,45 ± 0,33 |
| | ***ΔsidF,sidL***$^{X,V}$ | 0,096 ± 0,007 | 0,578 ± 0,007 | 0,674 ± 0,018 | ND | ND | 7,92 ± 0,49 | 3,51 ± 0,19 |
| | ***ΔsidF,sidL***$^{X,V,(AKL)}$ | 0,080 ± 0,004 | 0,459 ± 0,026 | 0,498 ± 0,057 | ND | ND | 6,59 ± 0,05 | 2,67 ± 0,09 |
| | ***ΔsidF*** | 0,110 ± 0,003 | 0,663 ± 0,004 | 0,679 ± 0,004 | ND | ND | 16,51 ± 0,03 | 4,12 ± 0,04 |

**Supplementary Table S5:** Absolute values (mean ± SD) of experiments presented in Figure **7**. The TAFC content of the supernatant and the FC content of the mycelia were normalized to mg biomass. ND, not detected.

| Strain | Biomass (g) -Fe | Biomass (g) +1 µm Fe | Biomass (g) +Fe | TAFC (nmol/mg) -Fe | TAFC (nmol/mg) +1 µm Fe | FC (nmol/mg) -Fe | FC (nmol/mg) +1 µm Fe |
|---|---|---|---|---|---|---|---|
| **wt** | 0,116 ± 0,002 | 0,491 ± 0,004 | 0,551 ± 0,012 | 38,83 ± 2,48 | 16,74 ± 0,25 | 4,13 ± 0,22 | 3,56 ± 0,04 |
| ***ΔsidF*** | 0,121 ± 0,004 | 0,483 ± 0,006 | 0,583 ± 0,013 | ND | ND | 16,79 ± 0,53 | 3,89 ± 0,08 |
| ***ΔsidL*** | 0,123 ± 0,004 | 0,484 ± 0,005 | 0,589 ± 0,016 | 41,28 ± 1,71 | 17,51 ± 0,27 | 1,14 ± 0,06 | 0,85 ± 0,02 |
| ***ΔpexC*** | 0,115 ± 0,003 | 0,491 ± 0,008 | 0,646 ± 0,005 | 20,64 ± 2,70 | 8,03 ± 0,40 | 17,45 ± 0,17 | 4,04 ± 0,04 |
| ***ΔpexCΔsidF*** | 0,119 ± 0,003 | 0,472 ± 0,008 | 0,640 ± 0,009 | ND | ND | 16,63 ± 0,19 | 4,05 ± 0,04 |
| ***ΔpexCΔsidL*** | 0,108 ± 0,005 | 0,448 ± 0,002 | 0,629 ± 0,004 | 21,02 ± 1,59 | 10,87 ± 0,27 | 10,16 ± 0,75 | 3,53 ± 0,03 |

**Supplementary Table S6:** Strains used in this study.

| Strain | Description | Reference |
|---|---|---|
| A1160+ | CEA10; *ΔakuB*$^{ku80}$*::pyrG*$^{-}$*zeo, pyrG*$^{-}$*::pyrG*$^{Af}$*; MAT1-1* | |
| *ΔsidF* | A1160+; *ΔsidF::hph* | This study |
| *ΔsidL* | A1160+; *ΔsidL::ptrA* | This study |
| *ΔsidLΔsidF* | A1160+; *ΔsidL::ptrA; ΔsidF:: hph* | This study |
| *ΔsidA* | A1160+; *ΔsidA::hph* | (13) |
| *ΔsidL,sidF*$^{X,V,Nterm(AKL)}$ | A1160+; *ΔsidLΔsidF; ΔfcyB::PxylP:Venus:SidF N-term(AKL)* | This study |
| *ΔsidL,sidF*$^{X,V,Cterm}$ | A1160+; *ΔsidLΔsidF; ΔfcyB::PxylP:Venus:SidF C-term* | This study |
| *ΔsidL,sidF*$^{X,V}$ | A1160+; *ΔsidLΔsidF; ΔfcyB::PxylP:Venus:SidF* | This study |
| *ΔsidL,sidF*$^{X,Nterm(AKL)}$ | A1160+; *ΔsidLΔsidF; ΔfcyB::PxylP:SidF N-term(AKL)* | This study |
| *ΔsidL,sidF*$^{X,Cterm(AKL)}$ | A1160+; *ΔsidLΔsidF; ΔfcyB::PxylP:SidF C-term* | This study |
| *ΔsidL,sidF*$^{X}$ | A1160+; *ΔsidLΔsidF; ΔfcyB::PxylP:SidF* | This study |
| *ΔsidF,sidL*$^{X,V}$ | A1160+; *ΔsidLΔsidF; ΔfcyB::PxylP:Venus:SidL* | This study |
| *ΔsidF,sidL*$^{X,V,(AKL)}$ | A1160+; *ΔsidLΔsidF; ΔfcyB::PxylP:Venus:SidL(AKL)* | This study |
| *ΔpexC* | A1160+; *ΔpexC::ptrA* | This study |
| *ΔpexCsidF* | A1160+; *ΔpexC::ptrA; ΔsidF::hph* | This study |
| *ΔpexCΔsidL* | A1160+; *ΔpexC::ptrA; ΔsidL::hph* | This study |
| *pexC*$^{C}$ | A1160+; *ΔpexC::ptrA; ΔfcyB::PpexC:pexC* | This study |

**Supplementary Table S7:** Primers used in this study.

| Plasmid | Description | 5'-3' Sequence |
|---|---|---|
| *sidF* | Fusion cassette (nested primers) | GAGCTGGGACGCAGAAAAGC |
| | | GGATATACAAAGGAAATGCATCCAGG |
| *sidL* | Fusion cassette (nested primers) | AAGCCGCCGTATGGACAGTA |
| | | ATCAAGCCGAACCAGGTTGT |
| | 5 UTR | GTCGTACAGACCTCATATGGCC |
| | | tagttctgttaccgagccggGGTGAGGGCGGACGGTCTAT |
| | *ptrA* | ccggctcggtaacagaactaACATCAGCAATTGATTACGGGATCCCATTGGT |
| | | gggagcatatcgttcagagcTATCTCCCTCTTGCATCTTTGTTTGTATTATA |
| | 3 UTR | gctctgaacgatatgctcccGCCTCATCCCGATTTCTTGG |
| | | TTTGTCTTTAGTATCTTGGCCAAGG |
| pSidF$^{X,V,}$ | Backbone | actcaagcttAAACAGCTATGACCATGATTAC |
| | | gcatcagtgcTGTGAAATTGTTATCCGCTC |
| | *xylP venus* | agcggataacaatttcacaGCACTGATGCGAGCAACAG |
| | | ggacccggacccggacccCTTGTACAGCTCGTCCATGC |
| | *sidF* | gggtccgggtccgggtccATGGCGACTCAAAGCAGCA |
| | | aatcatggtcatagctgtttACTGTTTGTTGAGGAATTAG |
| pSidF$^{X,V,N\text{-}term(AKL)}$ | Backbone | actcaagcttAAACAGCTATGACCATGATTAC |
| | | gcatcagtgcTGTGAAATTGTTATCCGCTC |
| | *xylP venus* | agcggataacaatttcacaGCACTGATGCGAGCAACAG |
| | | ggacccggacccggacccCTTGTACAGCTCGTCCATGC |
| | *sidF N-term* | gggtccgggtccgggtccATGGCGACTCAAAGCAGCA |
| | | gtaattacagctttgcatcccaACGTGGTCCAACCGGTGAGC |
| | 3 UTR | gctcaccggttggaccacgtTGGGATGCAAAGCTGTAATTAC |
| | | aatcatggtcatagctgtttACTGTTTGTTGAGGAATTAG |
| pSidF$^{X,V,C\text{-}term}$ | Backbone | actcaagcttAAACAGCTATGACCATGATTAC |
| | | gcatcagtgcTGTGAAATTGTTATCCGCTC |
| | *xylP venus* | agcggataacaatttcacaGCACTGATGCGAGCAACAG |
| | | ggacccggacccggacccCTTGTACAGCTCGTCCATGC |
| | *sidF C-term* | gggtccgggtccgggtccCCGATATGGGCCGTCGAC |
| | | aatcatggtcatagctgtttACTGTTTGTTGAGGAATTAG |
| pSidF$^{X}$ | Backbone | actcaagcttAAACAGCTATGACCATGATTAC |
| | | gcatcagtgcTGTGAAATTGTTATCCGCTC |
| | *xylP* | agcggataacaatttcacaGCACTGATGCGAGCAACAG |
| | | tgctgctttgagtcgccatGGTTGGTTCTTCGAGTCG |
| | *sidF* | gggtccgggtccgggtccATGGCGACTCAAAGCAGCA |
| | | aatcatggtcatagctgtttACTGTTTGTTGAGGAATTAG |
| pSidF$^{X,N\text{-}term(AKL)}$ | Backbone | actcaagcttAAACAGCTATGACCATGATTAC |
| | | gcatcagtgcTGTGAAATTGTTATCCGCTC |
| | *xylP* | agcggataacaatttcacaGCACTGATGCGAGCAACAG |
| | | tgctgctttgagtcgccatGGTTGGTTCTTCGAGTCG |
| | *sidF N-term* | gggtccgggtccgggtccATGGCGACTCAAAGCAGCA |
| | | gtaattacagctttgcatcccaACGTGGTCCAACCGGTGAGC |
| | 3 UTR | gctcaccggttggaccacgtTGGGATGCAAAGCTGTAATTAC |
| | | aatcatggtcatagctgtttACTGTTTGTTGAGGAATTAG |
| PSidF$^{X,C\text{-}term}$ | Backbone | actcaagcttAAACAGCTATGACCATGATTAC |
| | | gcatcagtgcTGTGAAATTGTTATCCGCTC |
| | *xylP* | agcggataacaatttcacaGCACTGATGCGAGCAACAG |
| | | gtcgacggcccatatcgccatGGTTGGTTCTTCGAGTCG |
| | *sidF C-term* | gggtccgggtccgggtccCCGATATGGGCCGTCGAC |
| | | aatcatggtcatagctgtttACTGTTTGTTGAGGAATTAG |

| | | |
|---|---|---|
| pSidL$^{X,V}$ | Backbone | actcaagcttAAACAGCTATGACCATGATTAC<br>gcatcagtgcTGTGAAATTGTTATCCGCTC |
| | *xylP venus* | agcggataacaatttcacaGCACTGATGCGAGCAACAG<br>ggacccggacccggacccCTTGTACAGCTCGTCCATGC |
| | *sidL* | gggtccgggtccgggtccATGCGTAACACTATCTGCCGT<br>aatcatggtcatagctgtttTTTGTCTTTAGTATCTTGGCCAAGG |
| pSidL$^{X,V,(AKL)}$ | Backbone | actcaagcttAAACAGCTATGACCATGATTAC<br>gcatcagtgcTGTGAAATTGTTATCCGCTC |
| | *xylP venus* | agcggataacaatttcacaGCACTGATGCGAGCAACAG<br>ggacccggacccggacccCTTGTACAGCTCGTCCATGC |
| | *sidL* | gggtccgggtccgggtccATGCGTAACACTATCTGCCGT<br>gtaattacagctttgcatcccaCTTCTCATTGTCCACCCGTG |
| | 3 UTR | cgggcctcgCTCGTCGGCGAGTCCAGC<br>aatcatggtcatagctgtttACTGTTTGTTGAGGAATTAG |
| *pexC* | Fusion cassette | TAGATGATAATAGGGCAAGCCGA<br>GACACAATCAGCAGTGACCC |
| | 5 UTR | TAGATGATAATAGGGCAAGCCGA<br>tagttctgttaccgagccggGATAGAGCTAGAGATGGGAGAGC |
| | *ptrA* | ccggctcggtaacagaactaACATCAGCAATTGATTACGGATCCCATTGGT<br>gggagcatatcgttcagagcTATCTCCCTCTTGCATCTTTGTTTGTATTATA |
| | 3 UTR | gctctgaacgatatgctcccGCTGCTTCAATTCCAGAGGG<br>GACACAATCAGCAGTGACCC |
| *sidL hph* | Fusion cassette<br>(nested primers) | AAGCCGCCGTATGGACAGTA<br>ATCAAGCCGAACCAGGTTGT |
| | 5 UTR | GTCGTACAGACCTCATATGGCC<br>tagttctgttaccgagccggGGTGAGGGCGGACGGTCTAT |
| | *hph* | ccggctcggtaacagaactaACGGCGTAACCAAAAGTCAC<br>gggagcatatcgttcagagcTCTTGACGACCGTTGATCTG |
| | 3 UTR | gctctgaacgatatgctcccGCCTCATCCCGATTTCTTGG<br>TTTGTCTTTAGTATCTTGGCCAAGG |
| pPexC$^{C}$ | Backbone | actcaagcttAAACAGCTATGACCATGATTAC<br>gcatcagtgcTGTGAAATTGTTATCCGCTC |
| | Native promoter/*pexC*/ 3 UTR | aatcatggtcatagctgtttCCTTTTAGAATATCCCTCAC<br>gagcggataacaatttcacaAATAAAAGGGATCTTCGGTG |

**Table S8:** Primers used for amplification of the digoxigenin-labelled probes for Northern- and Southern blot analysis.

| Probe | Gene | 5'-3' Sequence |
|---|---|---|
| *sidF* $^{N\text{-}term}$ | AFUA_3G03400 | CAAAGCAGCACCGAGCTTCC<br>ACGTGGTCCAACCGGTGAGC |
| *sidF* $^{C\text{-}term}$ | AFUA_3G03400 | ACCACGGGATTCACGGCCTG<br>CGCCATCCGATCCGAACTTT |
| *Venus* | Venus A206K | GACGTAAACGGCCACAAGTT<br>GAACTCCAGCAGGACCATGT |
| 3' NCR *fcyB* | AFUB_025700 | TGCGGTTTTTGGGTTTTATC<br>AGACCGTTGTTTCATACCGC |
| 5'NCR *sidF* | AFUA_3G03400 | CAATAACTGGTCATGCTCC<br>CGCGCAATGAGTTTG |
| 5'NCD *sidL* | AFUA_1G04450 | GTCGTACAGACCTCATATGGCC<br>tagttctgttaccgagccggGGTGAGGGCGGACGGTCTAT |
| 3'NCD *pexC* | AFUA_5G06300 | gctctgaacgatatgctcccGCTGCTTCAATTCCAGAGGG<br>GACACAATCAGCAGTGACCC |

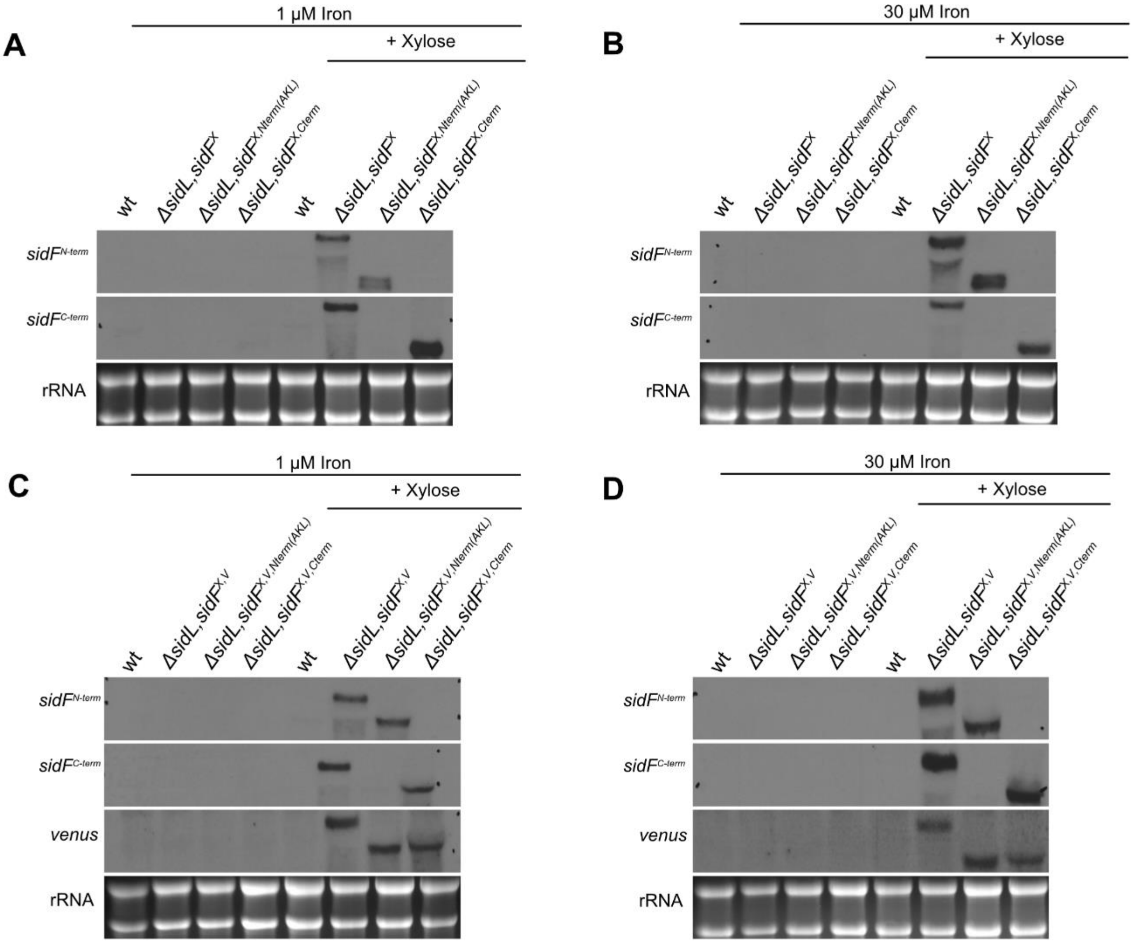


**Supplementary Figure S1: P*xylP*-mediated expression allows xylose-inducible production of full-length SidF (SidF$^{V}$), the N-terminal SidF domain (SidF$^{Nterm(AKL)}$, SidF$^{V,Nterm(AKL)}$) and the C-terminal SidF domain (SidF$^{Nterm}$, SidF$^{V,Nterm}$) with and without Venus-tag, independent of the iron availability, as shown at the transcript level.** Shake flask cultures were performed as described in Figure 3, without or with 1% xylose supplementation. (A) Venus-untagged alleles with 1 µM iron supplementation; (B) Venus-untagged alleles with 30 µM iron supplementation; (C) Venus-tagged alleles with 1 µM iron supplementation; (D) Venus-tagged alleles with 30 µM iron supplementation. rRNA is shown as a control for quality and loading of the RNA samples. Hybridization probes used were *sidF*$^{N\text{-}term}$ (specific for the gene sequence encoding the SidF N-terminal domain), *sidF*$^{C\text{-}term}$ (specific for the gene sequence encoding the SidF C-terminal domain) and *venus* (specific for the gene sequence encoding Venus).

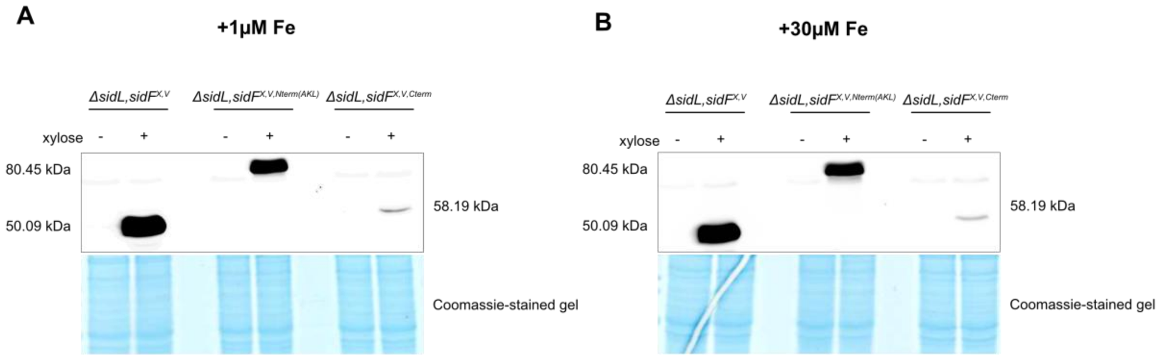


**Supplementary Figure S2: P*xylP*-mediated expression allows xylose-inducible production of full length SidF, the N-terminal SidF domain (SidF$^{V,Nterm(AKL)}$) and the C-terminal SidF domain (SidF$^{V,Nterm}$), independent of iron availability, as shown for Venus-tagged alleles at the protein level.** Shake flask cultures were performed as described in Figure 3. Respective strains were grown in shake flask cultures either with 1 µM (A) or 30 µM iron (B) supplementation, with and without xylose supplementation. Western blot analyses were performed with total cell extracts using a mouse α-GFP antibody. Coomassie-stained protein gels are shown as control for quality and loading of the protein samples. Notably, the C-terminal SidF domain showed significantly reduced protein levels compared to both full-length SidF and the N-terminal SidF domain, indicating reduced protein stability.

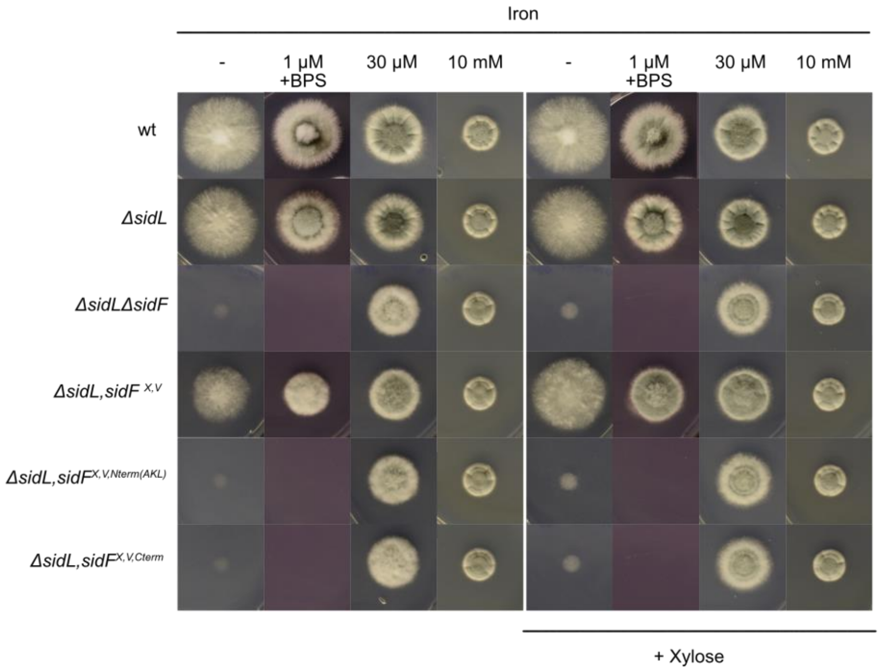


**Supplementary Figure S3: Individual expression of the Venus-tagged version of either the N-terminal (*ΔsidL,sidF*$^{X,V,Nterm(AKL)}$) or the C-terminal (*ΔsidL,sidF*$^{X,V,Cterm}$) SidF domains causes a growth defect similar *ΔsidFΔsidL* under iron limitation.** Experimental details are described in Figure 2. Cultures were either not supplemented or supplemented with 1 % xylose for P*xylP* induction.

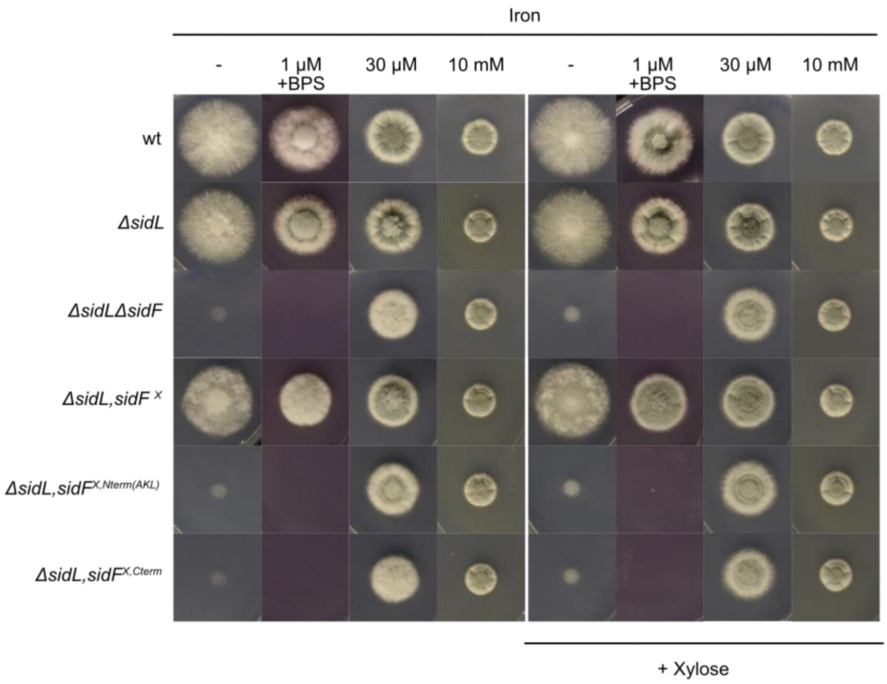


**Supplementary Figure S4: Individual expression of either the N-terminal (*ΔsidL,sidF*$^{X,V,Nterm(AKL)}$) or the C-terminal (*ΔsidL,sidF*$^{X,V,Cterm}$) SidF domains causes growth defects similar to *ΔsidFΔsidL* under iron limitation.** Experimental details are described in Figure 2. Cultures were either not supplemented or supplemented with 1 % xylose for P*xylP* induction.

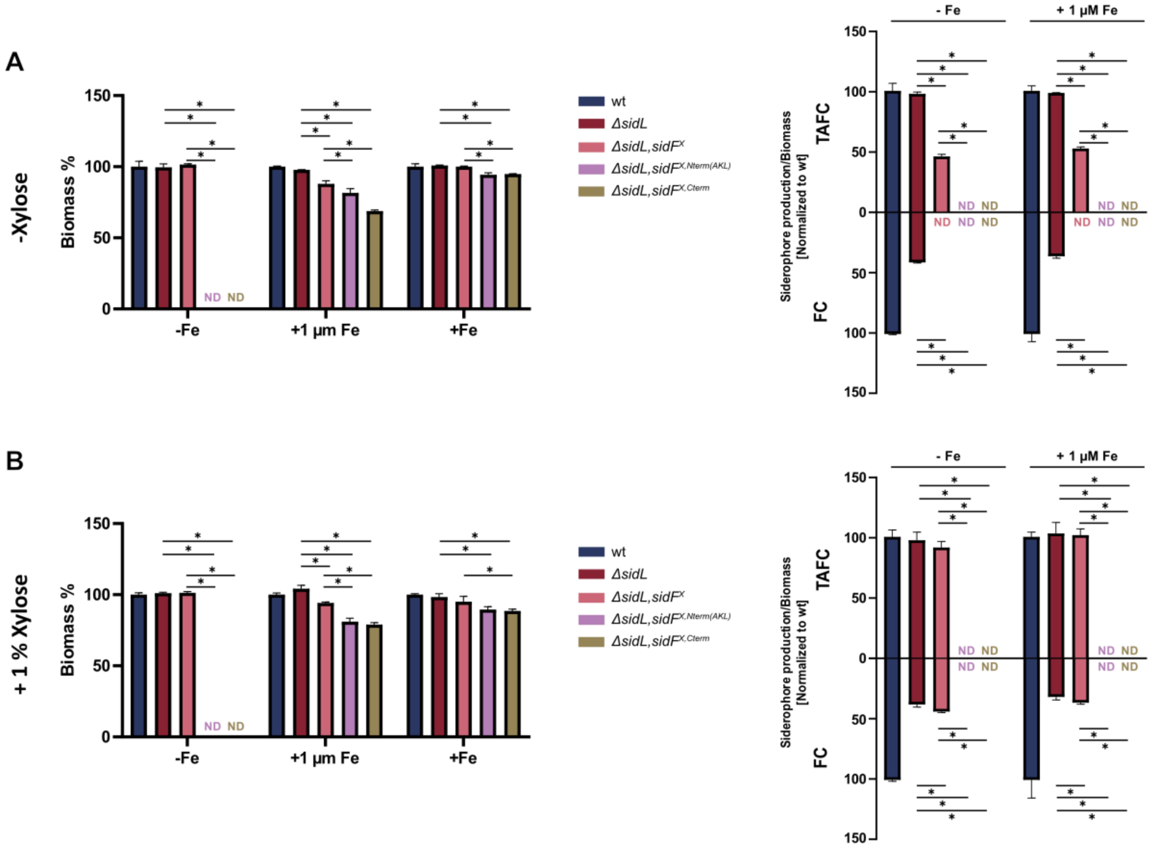


**Supplementary Figure S5: Neither the individual N- nor the C-terminal domain of SidF, without the Venus tag, supports biosynthesis of FusTS or FchTS *in vivo*.** Liquid shake flask cultures and determination of siderophore production were done as described in Figure 3. However, cultures were either not supplemented (A) or supplemented with 1 % xylose for P*xylP* induction (B). Statistically significant differences were calculated by one-way ANOVA ($p \leq 0.005$) compared to *ΔsidL* and *ΔsidL,sidF*$^{X,V}$, respectively, and are indicated with *. Absolute values are given in Supplementary Table S3.

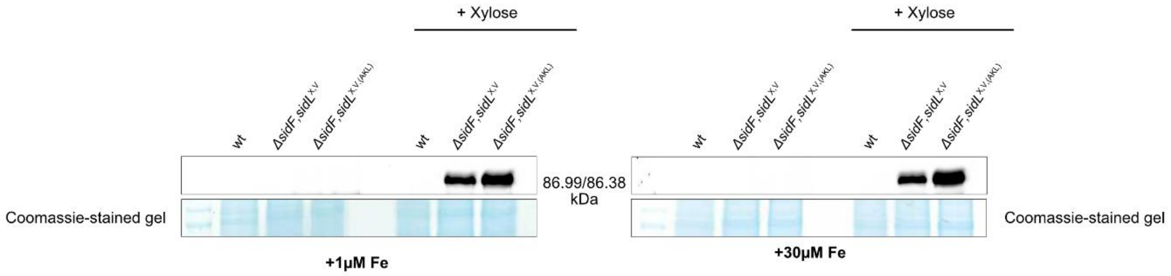


**Supplementary Figure S6: P*xylP*-mediated expression allows xylose-inducible production of N-terminal Venus tagged SidL without (*ΔsidF,sidL*$^{X,V}$) and with (*ΔsidF,sidL*$^{X,V,(AKL)}$) artificial C-terminal tagging with a PTS1 sequence, independent of iron availability.** Shake flask cultures were prepared as described in Figure 3. However, cultures were either not supplemented or supplemented with 1 % xylose for P*xylP* promoter induction. Western blot analyses were performed with total cell extracts using a mouse α-GFP antibody. Coomassie-stained protein gels are shown as control for quality and loading of the protein samples.

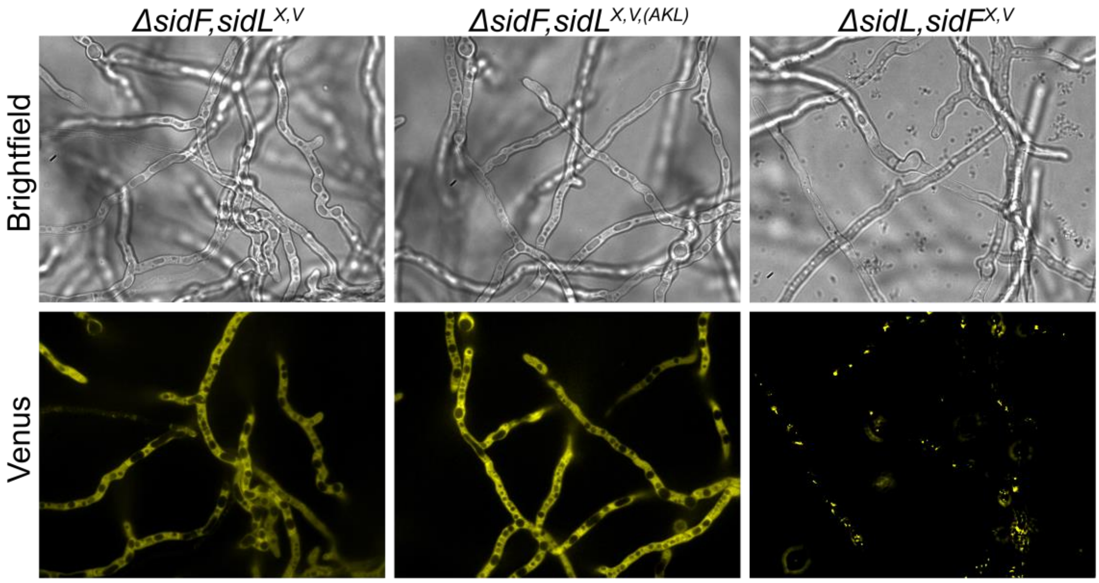


**Supplementary Figure S7: C-terminal tagging with an artificial PTS1 sequence (*ΔsidF,sidL*$^{X,V,(AKL)}$) does not target N-terminally Venus-tagged SidL to peroxisomes.** Fungal strains were grown for 16 h at 37 °C in 8-well chamber slides with 200 µl of minimal medium, supplemented with 30 µM iron and 0.1 % xylose, using an inoculum of $10^4$ spores per well. Strain *ΔsidL,sidF*$^{X,V}$, producing Venus-tagged SidL, is shown as example for a peroxisomal localized protein.

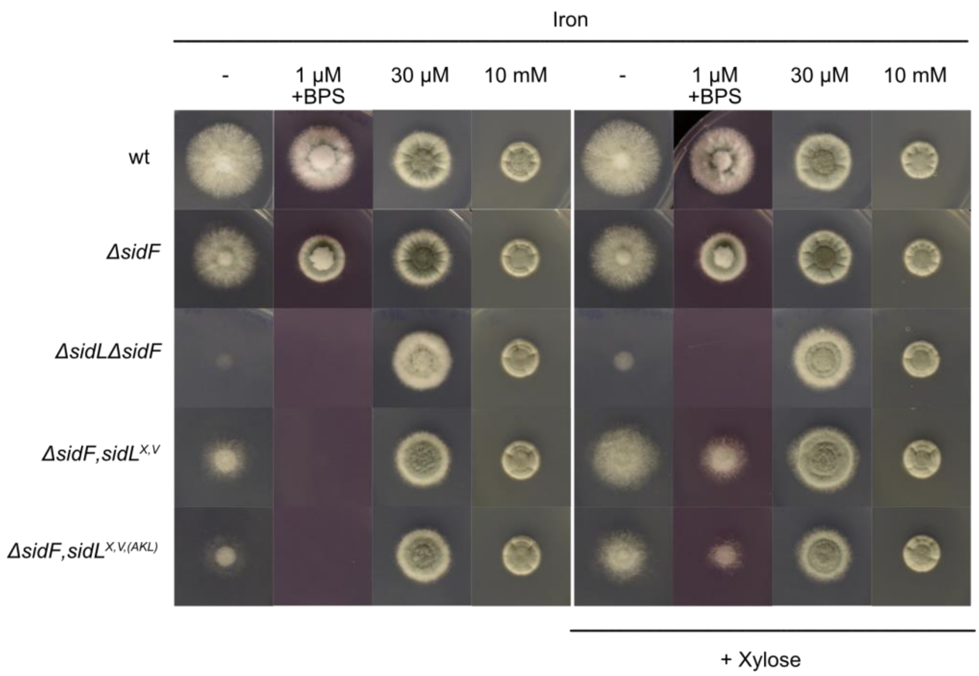


**Supplementary Figure S8: C-terminal PTS1 sequence tagging (*ΔsidF,sidL*$^{X,V,(AKL)}$) impairs the function of P*xylP*-controlled and N-terminally Venus-tagged SidL (*ΔsidF,sidL*$^{X,V}$).** Experimental details are described in Figure 2. Cultures were either not supplemented or supplemented with 1 % xylose for P*xylP* induction.

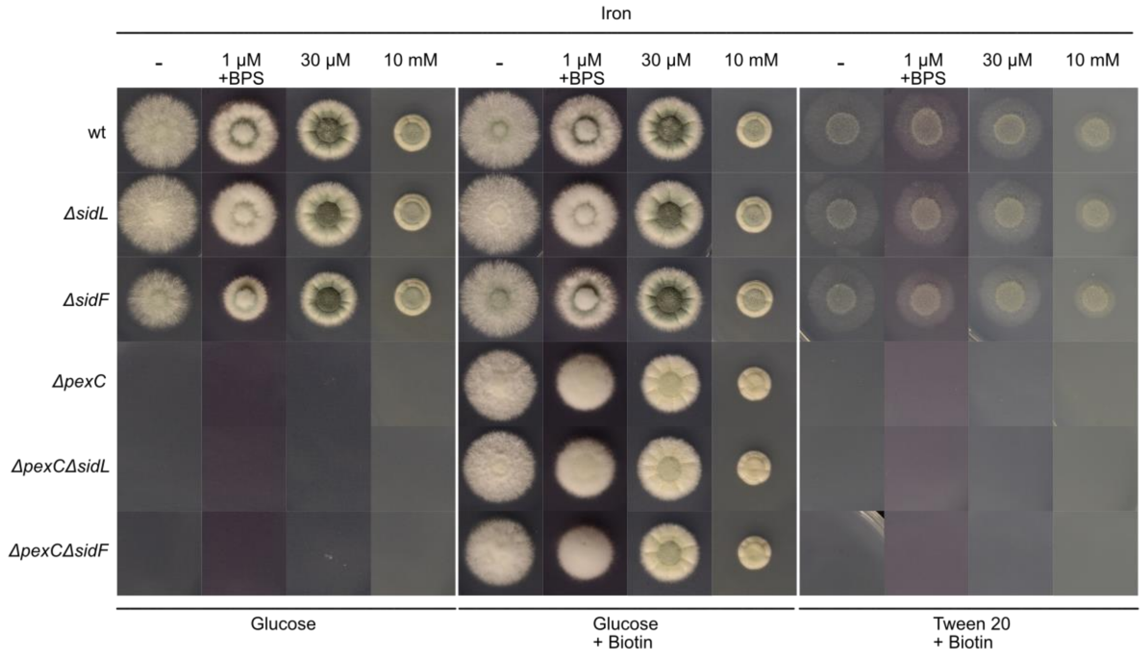


**Supplementary Figure S9: Lack of PexC blocks the utilization of fatty acids as carbon source, decreases conidial pigmentation and causes biotin auxotrophy.** Cultures were inoculated on media containing either 1 % glucose or 0.1 % Tween 20 (which contains oleic acid as the primary fatty acid) as sole carbon sources and were supplemented or not with biotin. In this growth assay, ammonium chloride was used as the sole nitrogen source instead of glutamine, as glutamine might also serve as a carbon source. Experimental details are described in Figure 2.

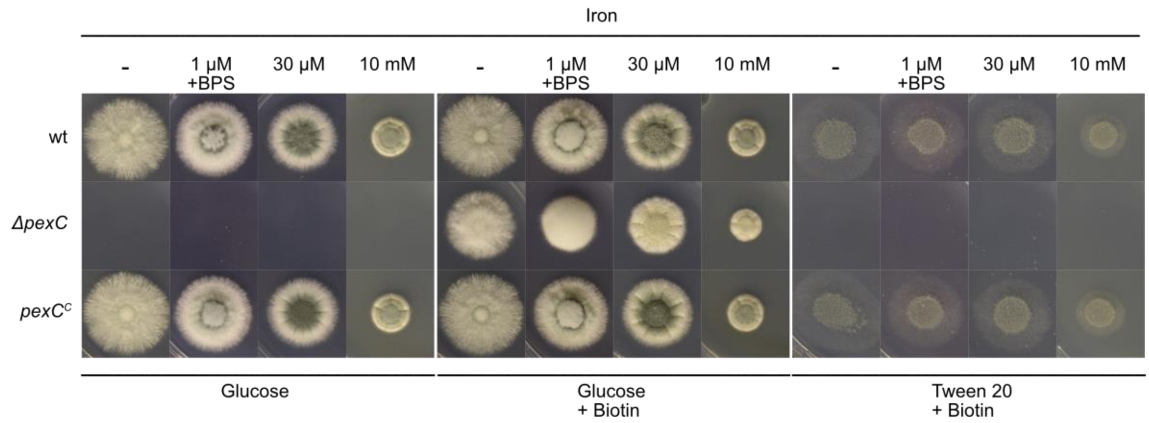


**Supplementary Figure S10: Reintegration of *pexC* cures the *ΔpexC* defect in utilization of fatty acids as sole carbon source, the decreased conidial pigmentation and the biotin auxotrophy.** Cultures were inoculated on media containing either 1 % glucose or 0.1 % Tween 20 (which contains oleic acid as the primary fatty acid) as sole carbon sources and were supplemented or not with biotin. In this growth assay, ammonium chloride was used as the sole nitrogen source instead of glutamine, as glutamine might also serve as a carbon source. Experimental details are described in Figure 2.

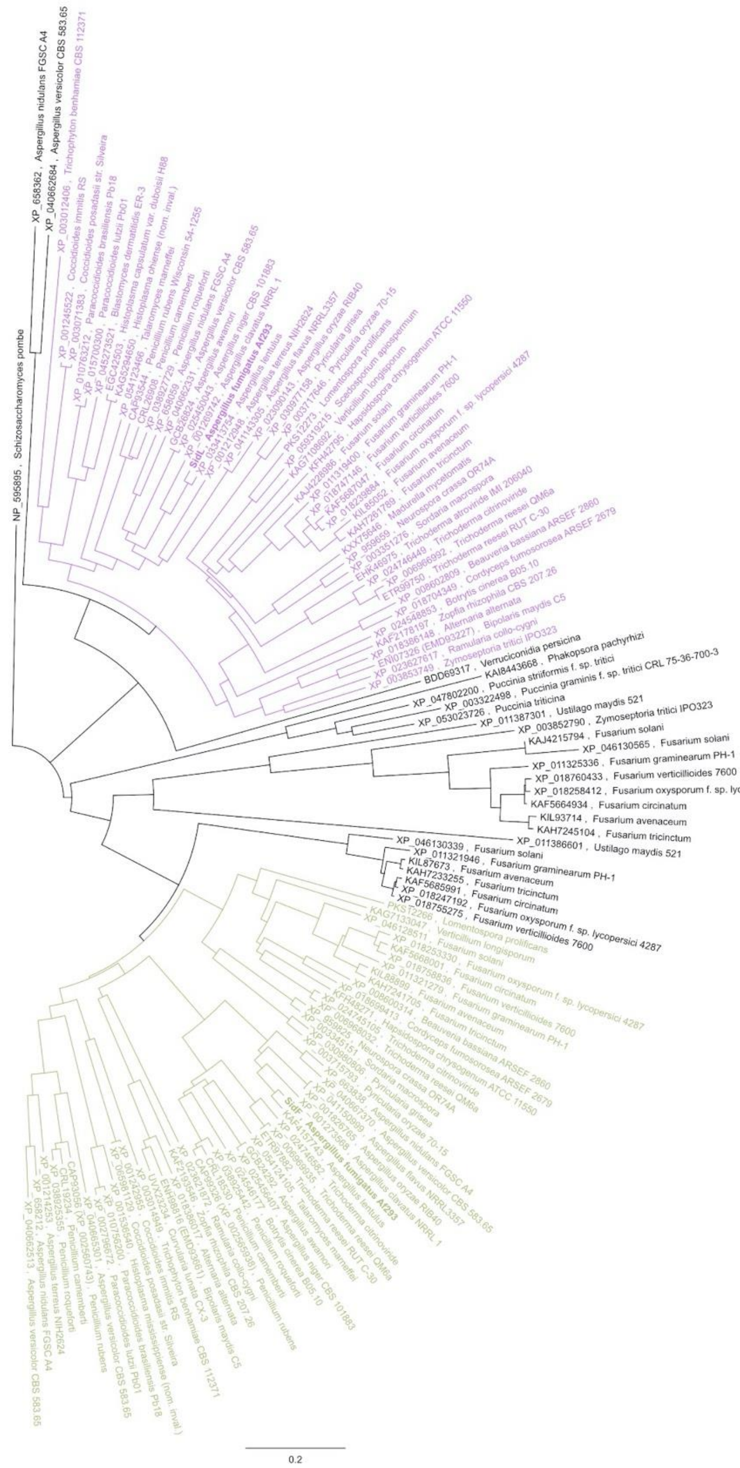


**Supplementary Figure S11: *A. fumigatus* SidF and SidL represent members of two phylogenetically separate clades, termed SidF and SidL clade, respectively.** SidL and SidF clades are colored in violet in green, respectively. *A. fumigatus* homologs are highlighted in bold. The phylogenetic analysis was generated using Geneious Prime and is based on NCBI blastp searches (51) including 82 fungal species; given are species names and accession numbers. Homologs were exclusively identified in hydroxamate-class siderophore producing fungi. Homologs were not found in *Batrachochytrium dendrobatidis*, *Candida albicans*, *Candida auris*, *Candida parapsilosis*, *Candida tropicalis*, *Colletotrichum coccodes*, *Cryptococcus gattii*, *Cryptococcus neoformans*, *Komagataella phaffii*, *Lacazia loboi*, *Mucor circinelloides*, *Mucor mucedo*, *Nakaseomyces glabratus (Candida glabrata)*, *Phanerochaete chrysosporium*, *Phycomyces blakesleeanus*, *Pichia kudriavzevii*, *Pneumocystis carinii*, *Pneumocystis jirovecii*, *Postia placenta*, *Rhizopus arrhizus*, *Rhizopus delemar*, *Rhizopus microspores*, *Saccharomyces cerevisiae*, and *Yarrowia lipolytica*

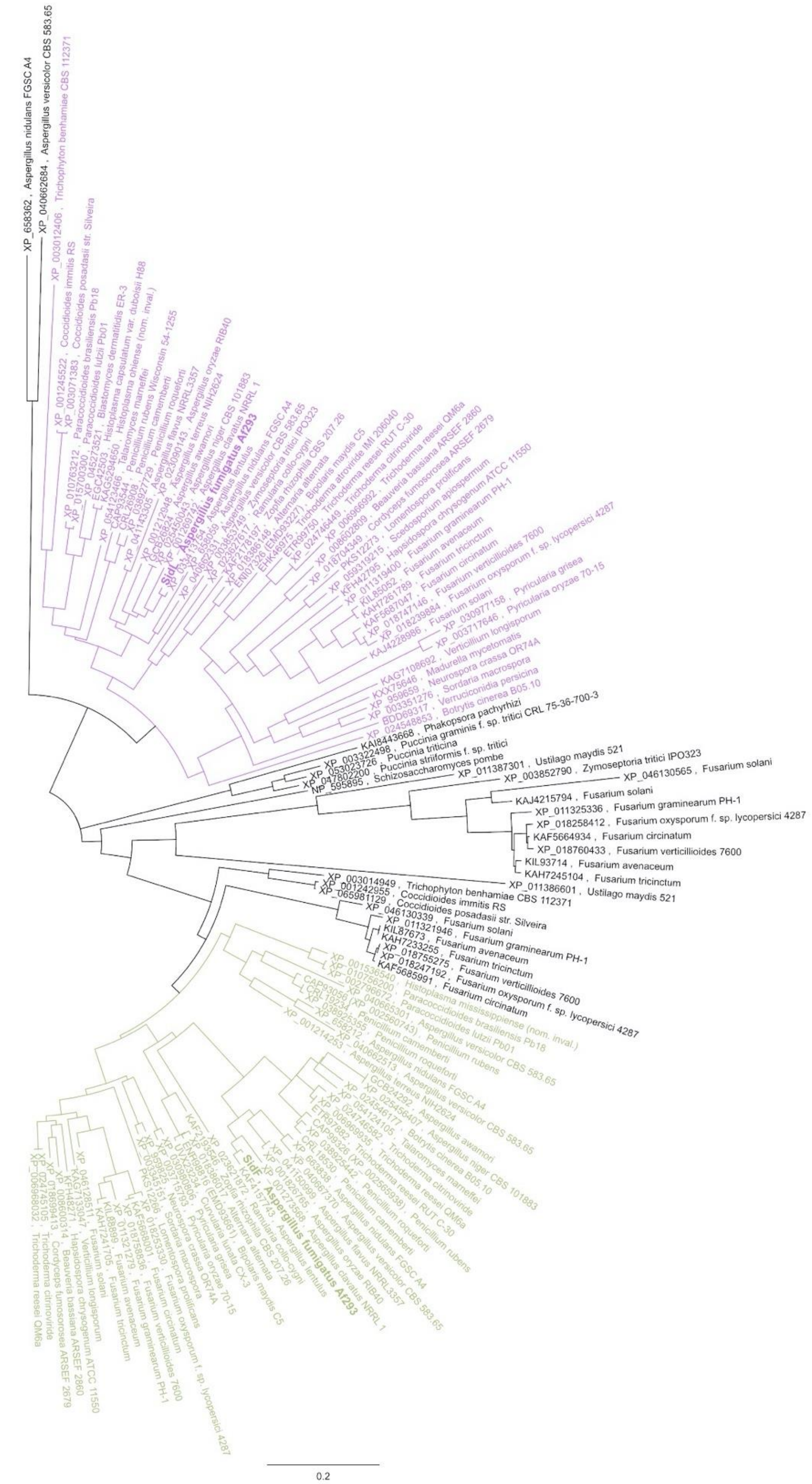


**Supplementary Figure S12:** The GNAT domains of SidF and SidL homologs cluster into two phylogenetically separate clades. SidL and SidF clades are colored in violet in green, respectively. *A. fumigatus* homologs are highlighted in bold. This phylogenetic analysis is based on the phylogenetic analysis of SidF and SidL homologs shown in *Figure S11*, but included only the GNAT domains extracted from the multiple alignment of all sequences using Geneious Prime.

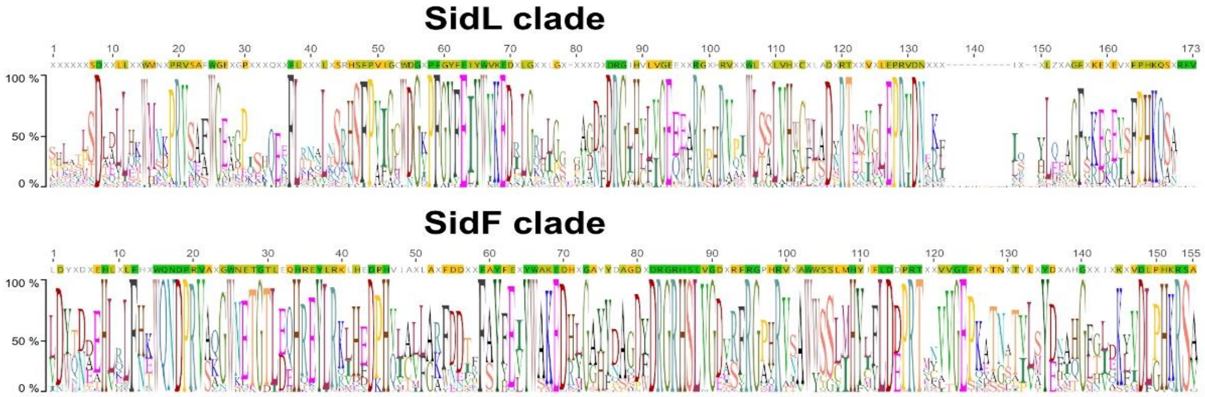


**Supplementary Figure S13: The GNAT domains of SidF and SidL clades show overlapping but different consensus sequences.** *S*equence logos of SidF and SidL clades were generated using Geneious Prime based on the phylogenetic analysis shown in *Figure S12*. Using the score matrix Blosum62, amino acid residues with 100% conservation are highlighted in green, with 80-99% in gold and with 60-79% in yellow.

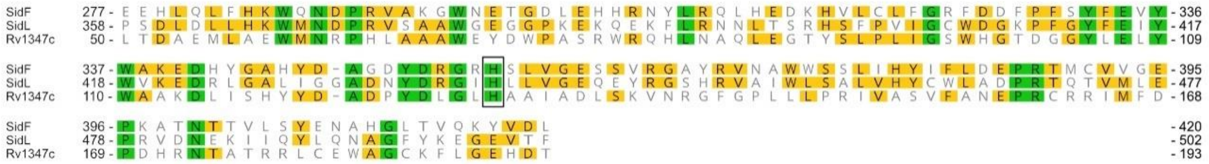


**Supplementary Figure S14: *Mutliple alignment of the GNAT-domains of M. tuberculosis Rv1347c and A. fumigatus SidF and SidL.*** Using the score matrix Blosum62, amino acid residues with 100% conservation are highlighted in green and with 60-79% in yellow. The histidine residue shown to be essential for activity of siderophore biosynthetic N$^6$-hydroxylysine acetyltransferase Rv1347c is boxed (38).

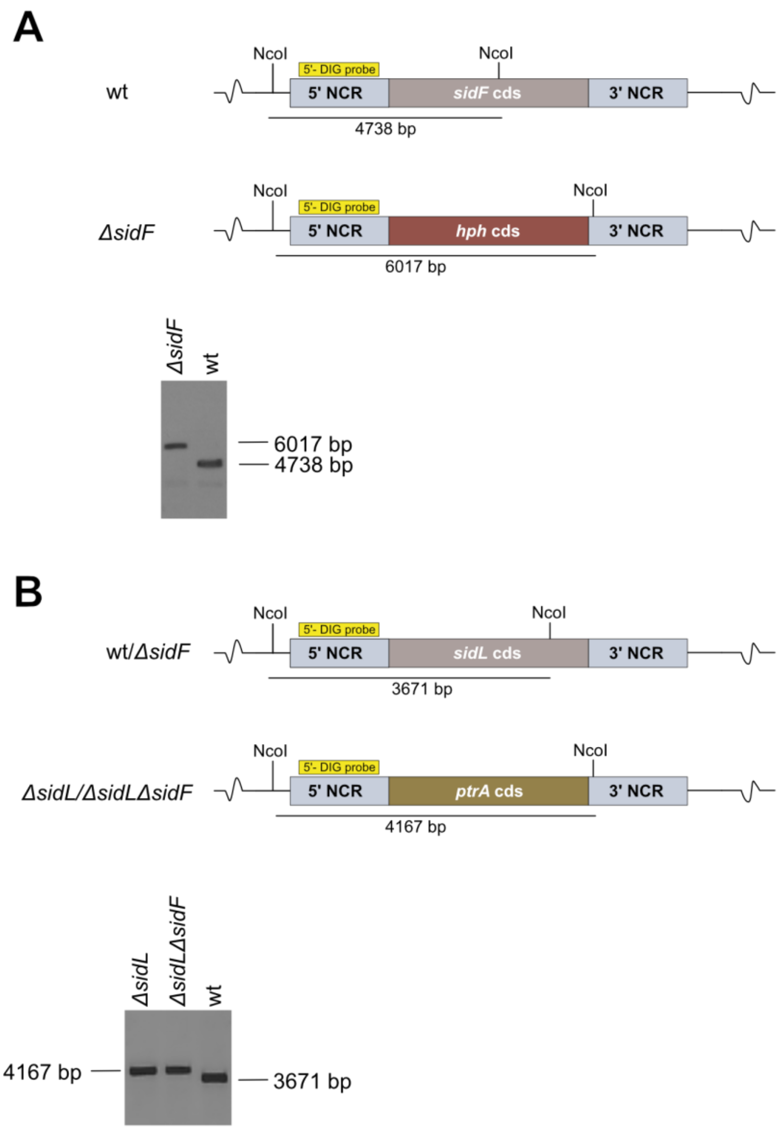


**Supplementary Figure S15: Scheme of the genomic loci for deletion of *sidF* and *sidL* in *A. fumigatus* and Southern blot verification of generated mutant strains.** Digestion using the restriction enzyme *Nco*I resulted in a 6,017 bp fragment in wt and in a 4,738 bp fragment in *ΔsidF* (A). Digestion using the restriction enzyme *Nco*I resulted in a 4,167 bp fragment in wt and *ΔsidF*, and a 3,671 bp fragment in *ΔsidL* and *ΔsidLΔsidF* (B). The hybridization probes indicated in (A) (5'NCD *sidF*) and (B) (5'NCD *sidL*) were generated from gDNA using primers mentioned in Supplementary Table S8. NCR, non-coding region.

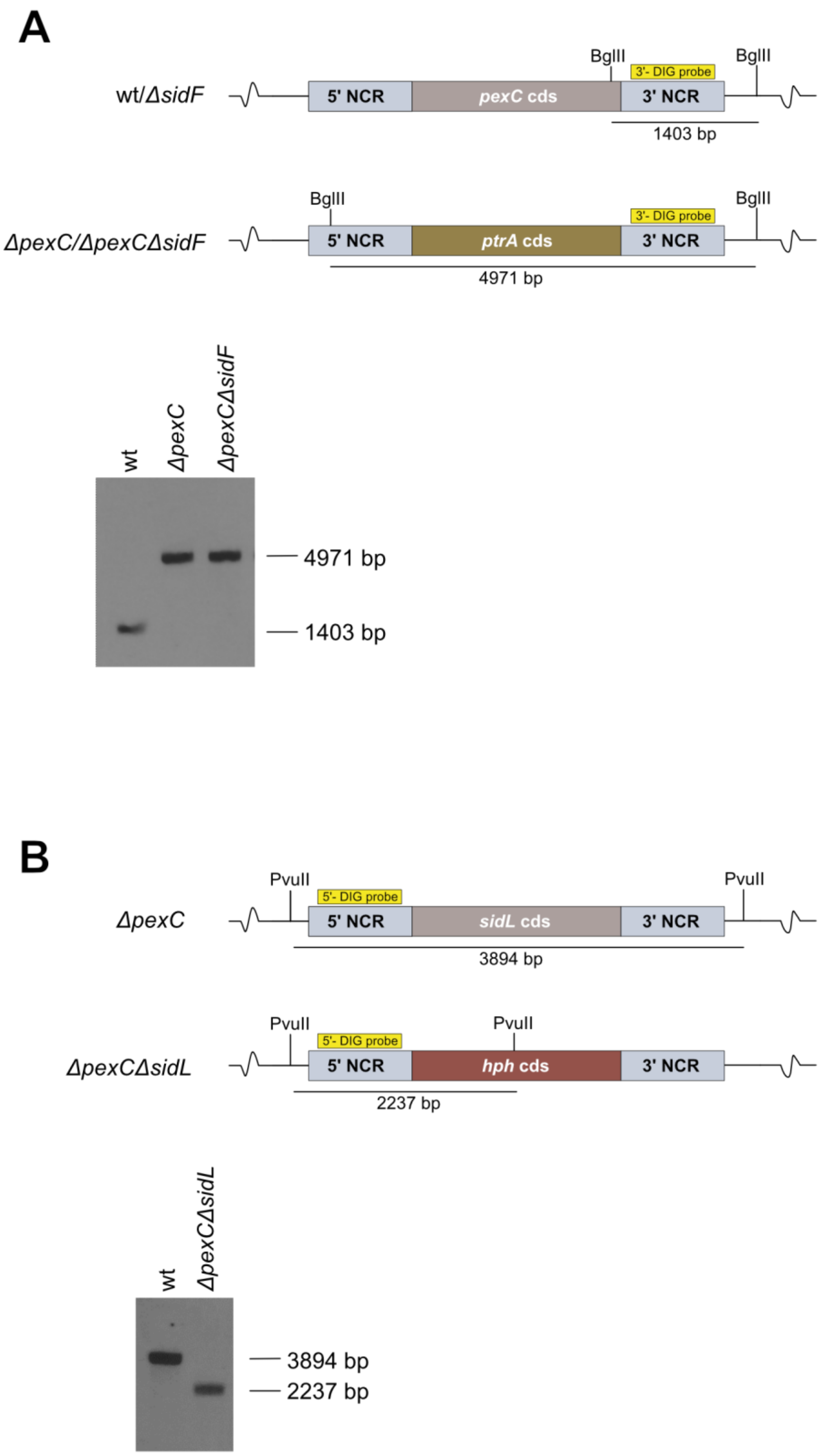


**Supplementary Figure S16: Scheme for the deletion of *pexC* in *A. fumigatus* and Southern blot verification of the generated mutant strain.** Digestion using the restriction enzyme *Bgl*II resulted in a 1,403 bp fragment in wt and a 4,931 bp fragment in *ΔpexC* and *ΔpexCΔsidF* (A). Scheme for deletion of *sidL* in *A. fumigatus ΔpexC* and Southern blot verification. Restriction digestion using the restriction enzyme *Pvu*II results in a 3,894 bp fragment in wt and a 2,237 bp fragment in *ΔpexCΔsidL* (B). The hybridization probes indicated in (A) (3'NCD *pexC*) and (B) (5'NCD *sidL*) were generated from gDNA using primers mentioned in Supplementary Table S8. NCR, non-coding region.

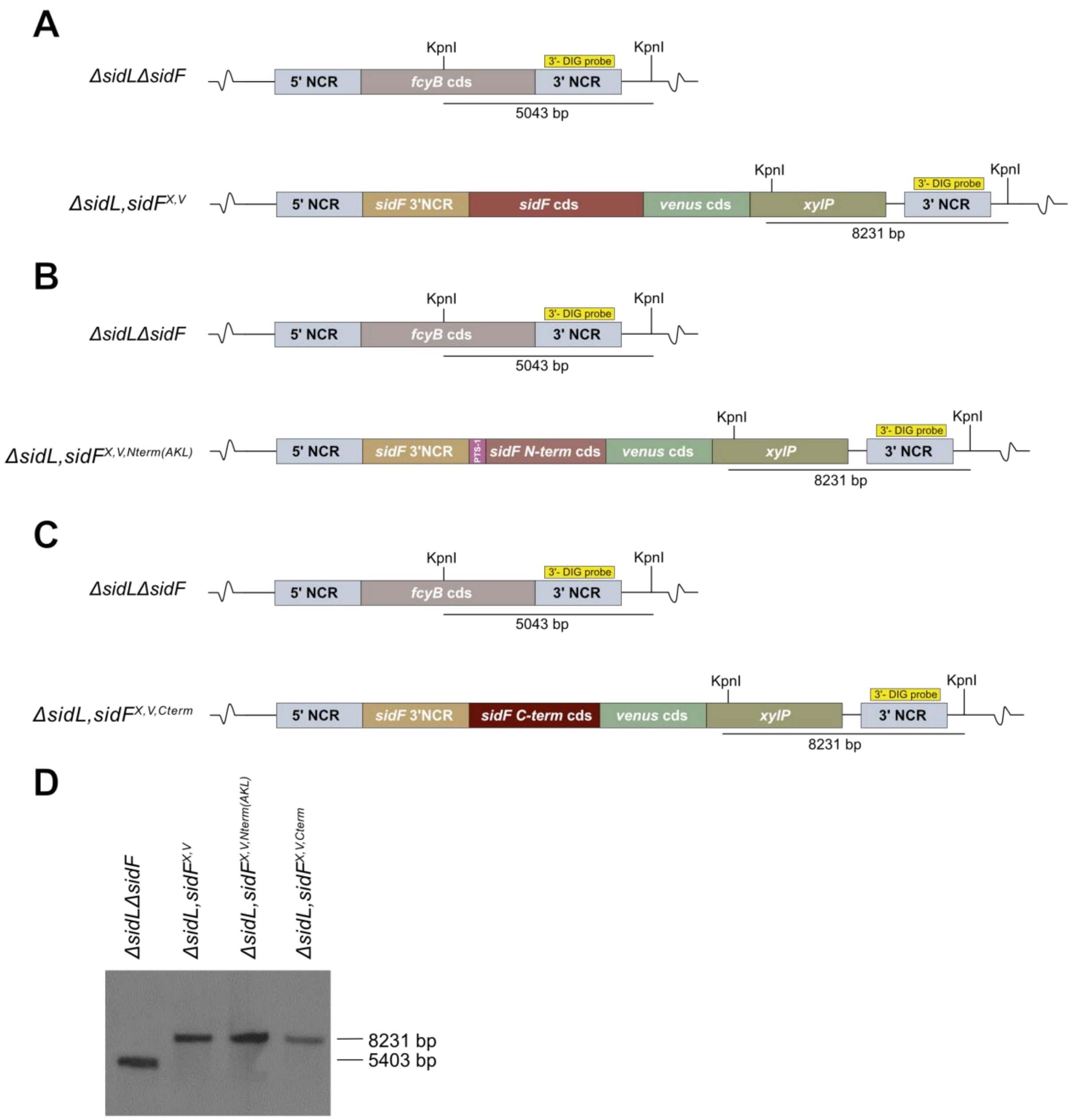


**Supplementary Figure S17: Scheme for the generation of the *ΔsidL sidF*$^{X,V}$ (A), *ΔsidL sidF*$^{X,V,Nterm(AKL)}$ (B) and *ΔsidL sidF*$^{X,V,Cterm}$ (C) strains in *A. fumigatus* and verification of generated mutant strains by Southern blot analysis (D).** Digestion using the restriction enzyme *Kpn*I resulted in a 5,403 bp fragment in *ΔsidLΔsidF* and an 8,231 bp fragment in the mutant strains. The hybridization probe indicated in (A), (B) and (C) (3'NCD *fcyB*) was generated from gDNA using primers mentioned in Supplementary Table S8. NCR, non-coding region.

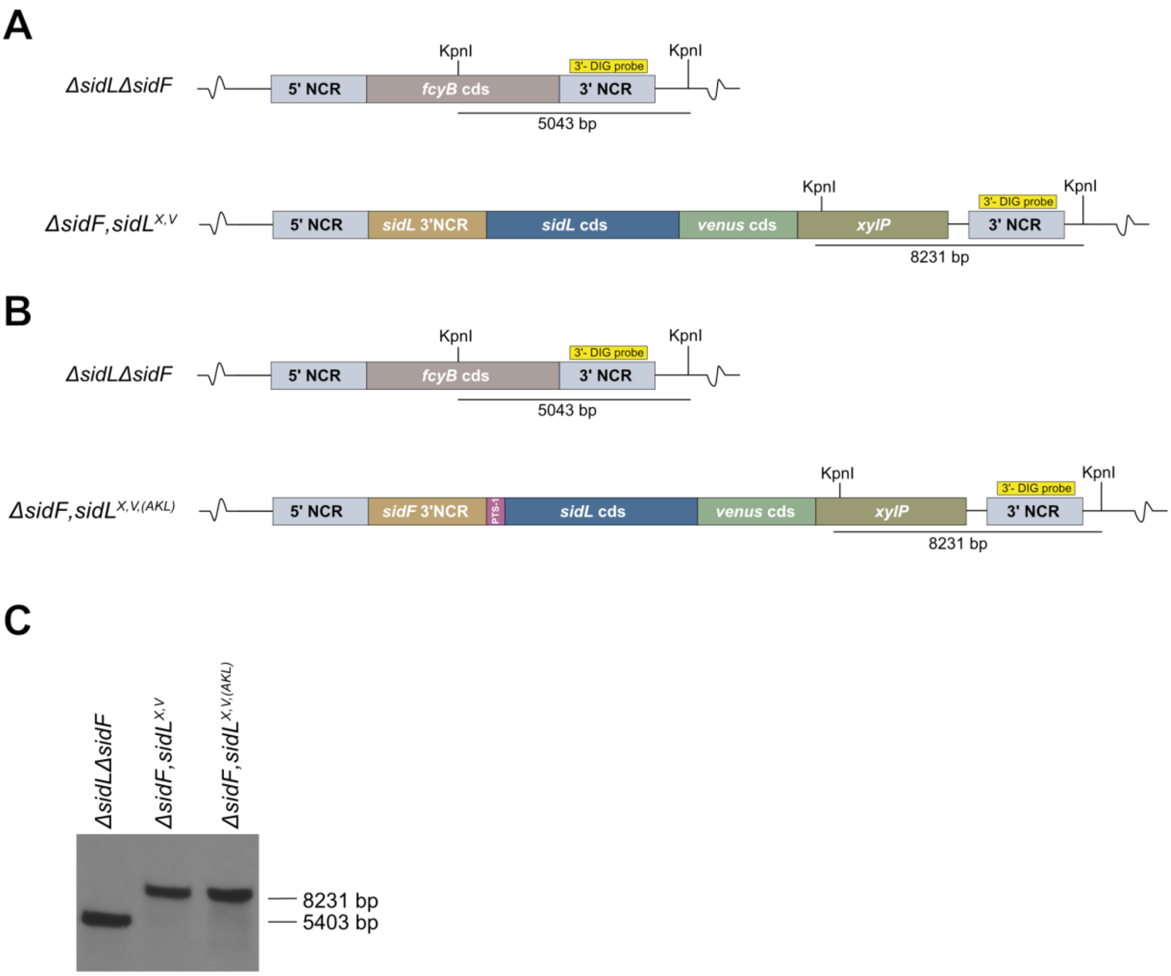


**Supplementary Figure S18: Scheme for the generation of the *ΔsidF sidL*$^{X,V}$ (A) and *ΔsidF sidL*$^{X,V,(AKL)}$ (B) strains in *A. fumigatus* and verification of generated mutant strains by Southern blot analysis (C).** Digestion using the restriction enzyme *Kpn*I resulted in a 5,403 bp fragment in *ΔsidLΔsidF* and an 8,231 bp fragment in the mutant strains. The hybridization probe labelled in A and B (3'NCD *fcyB*) was generated from gDNA using primers mentioned in Supplementary Table S8. NCR, non-coding region.

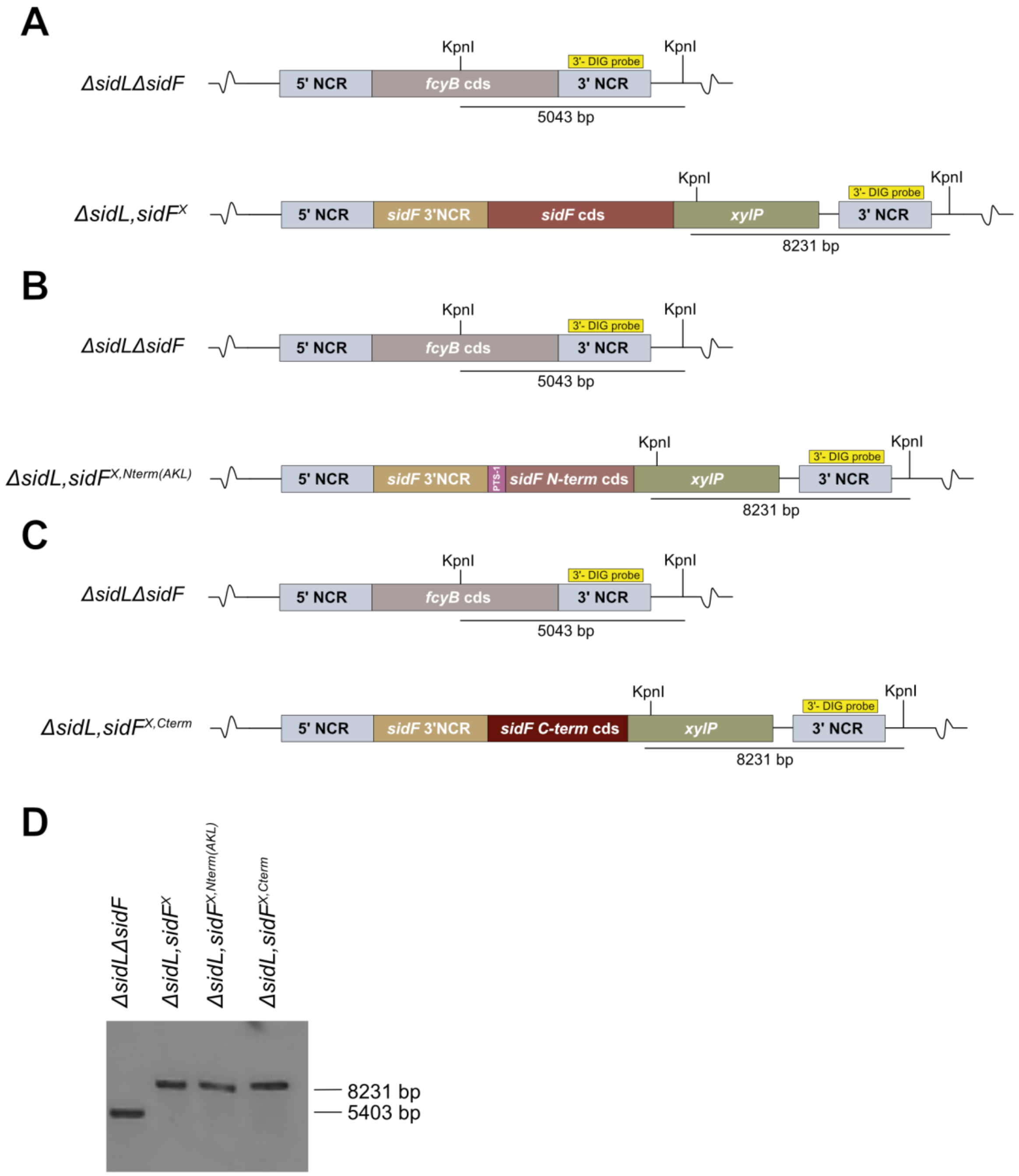


**Supplementary Figure S19: Scheme for the generation of the *ΔsidL sidF*$^{X}$ (A), *ΔsidL sidF*$^{X,Nterm(AKL)}$ (B) and *ΔsidL sidF*$^{X,Cterm}$ (C) strains in *A. fumigatus* and verification of generated mutant strains by Southern blot analysis (D).** Digestion using the restriction enzyme *Kpn*I resulted in a 5,403 bp fragment in *ΔsidLΔsidF* and an 8,231 bp fragment in the mutant strains. The hybridization probe indicated in (A), (B) and (C) (3'NCD *fcyB*) was generated from gDNA using primers mentioned in Supplementary Table S8. NCR, non-coding region.

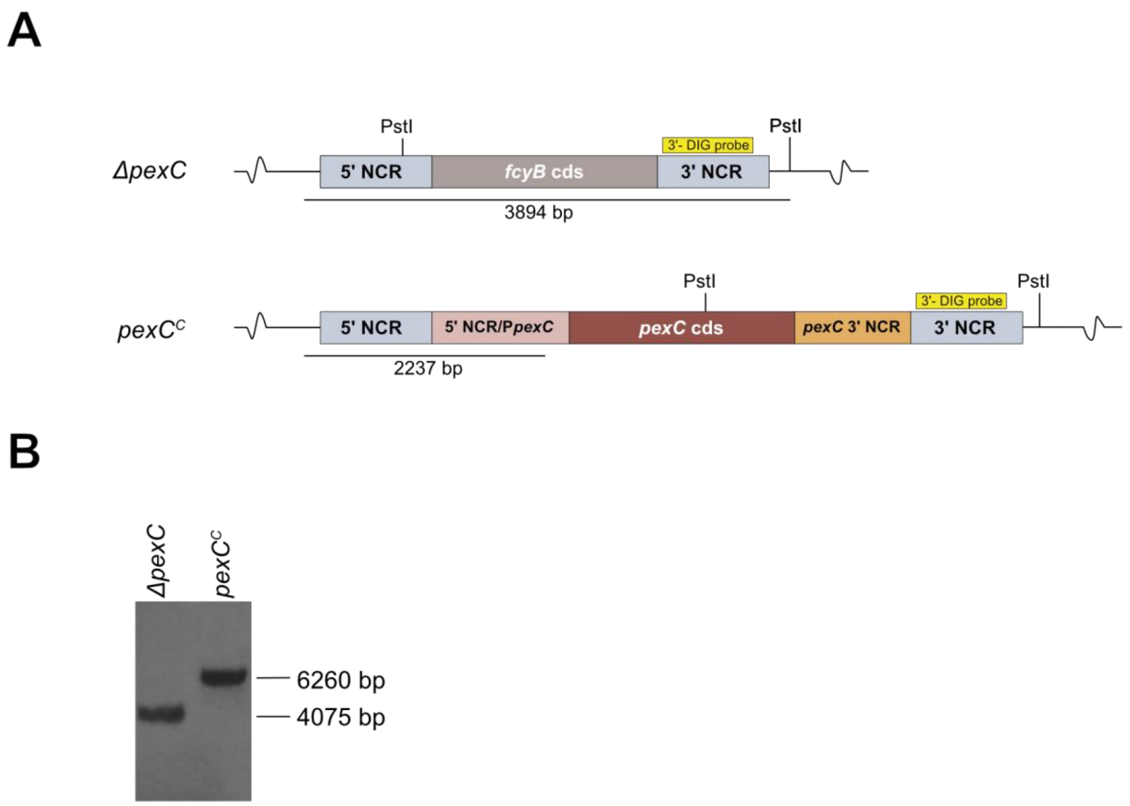


**Supplementary Figure S20: Scheme for the generation of the *pexC*$^{Comp}$ strain in *A. fumigatus* (A) and results of the Southern blot analysis for verification of the mutant strain (B).** Digestion using the restriction enzyme *Pst*I resulted in a 4,075 bp fragment in *ΔpexC* and a 6,260 bp fragment in the mutant strain. The hybridization probe indicated in (A) (3'NCD *fcyB*) was generated from gDNA using primers mentioned in Supplementary Table S8. NCR, non-coding region.
